# The representation of emotion knowledge in hippocampal-prefrontal systems

**DOI:** 10.1101/2025.06.03.657722

**Authors:** Yumeng Ma, Philip A. Kragel

## Abstract

Emotional experiences involve more than bodily reactions and momentary feelings—they depend on knowledge about the world that spans contexts and time. Although it is well established that individuals conceptualize emotions using a low-dimensional space organized by valence and arousal, the neural mechanisms giving rise to this configuration remain unclear. Here, we examine whether hippocampal-prefrontal circuits—regions implicated in forming cognitive maps—also support the abstraction of emotional experiences. Using functional MRI data collected as participants viewed emotionally evocative film clips, we found that activity in hippocampal and prefrontal cortex predicted self-reported emotion across schematically distinct videos, consistent with a role in structural learning. Computational modeling of emotion transitions revealed that hippocampal responses to films and emotion self-reports could be predicted based on the statistical regularities of emotion transitions across different temporal scales. These findings demonstrate that hippocampal-prefrontal systems represent emotion concepts at multiple levels of abstraction, offering new insight into how the brain organizes emotion knowledge.

## Main Text

Our experience of emotions is more than just momentary reactions to the world and accompanying feelings. Humans rely on emotion concepts—knowledge that helps us categorize, communicate, and make sense of emotional events in our lives. Some emotion knowledge is grounded in the particulars of individual episodes, such as the adorable features of a specific childhood pet or the growl of an aggressive neighborhood dog. Other knowledge concerns properties of events that generalize across episodes, like an opportunity for reward or the presence of a threat. Even though emotional events can differ widely in what we see, hear, think, or feel, we learn to abstract them into distinct categories of variable yet related instances.

Behavioral evidence suggests that humans compress information about complex emotional episodes into a low-dimensional affective space^1–4^. When individuals judge the meaning of words^5,6^, emotions conveyed by others^7^, predict how others are likely to feel in a given situation^8^, or self-report on their own experience^9^, their ratings primarily vary along dimensions of valence and arousal, suggesting that these variables are the most fundamental property of the mind^3^. Based on these observations, it has been suggested that humans represent the structure of affect in a map-like way^3,5^, using dimensions of valence and arousal as the organizing axes. If represented in such a format, emotion concepts would be positioned at specific locations in a relational network, much like landmarks are placed on a Cartesian map. With experience, individuals would learn the structure of affective space, enabling them to predict transitions from one experience to the next (e.g., the tendency to shift from a state of anxiety to fear as a threat becomes more proximal^10^) and to make decisions based on the anticipated consequences of actions^11^.

Neuroscience research has begun to characterize how the brain represents knowledge about emotional events. Work in nonhuman animals^12–16^ and human neuroimaging studies^17–21^ has identified functionally dissociable systems involved in valence and arousal processing. Neuroimaging has revealed that different categories of emotional events are differentiated by patterns of activity that are distributed across subcortical and cortical brain networks^22–24^, with the content represented varying across regions. Responses to emotional situations range from more situation-dependent activity in sensory cortices^25–28^ and more general category representations in transmodal cortical areas^29,30^. Together, these studies show that the brain represents emotional events using multiple systems in parallel, and that information processing occurs in a hierarchal fashion.

Although it is well-established that multiple aspects of emotional events are reflected across multiple brain systems, it remains unclear how emotionally relevant signals are transformed into low-dimensional affective spaces. One possibility is that memory systems necessary for organizing knowledge across domains are involved in mapping relationships between emotional events. Evidence suggests that the hippocampal formation—known for its involvement in spatial navigation^31^, memory^32^, and emotion^33–35^—represents knowledge in a map-like way^36–40^. To build a map of the environment, neural populations in the hippocampus bind together highly processed sensory inputs to form sparse representations of specific locations^31^ and hierarchically organized concepts^41^. Based on the coactivation of multiple concepts, neural populations in the entorhinal cortex^42^ and ventromedial prefrontal cortex (vmPFC) learn relations among concepts using a grid-like code^36–40^. If similar mechanisms are used to construct maps of emotion concepts^43^, then knowledge about distinct sets of emotional experiences should be encoded in patterns of hippocampal activity, whereas the spatial bases that define affective space (i.e., grid-like codes that span dimensions of valence and arousal) should be present in entorhinal cortex and vmPFC^44^.

Here we evaluate the proposal^43^ that hippocampal-prefrontal systems represent emotional events in a map-like way. We analyze fMRI signals in hippocampal-prefrontal systems acquired as human participants watched a series of cinematic videos (sampled from the Emo-FilM dataset^45^). As information about prototypical emotional events is typically accessible to consciousness, we first probe whether self-report measures of emotional experience can be decoded from patterns of hippocampal-prefrontal activity, and whether representations of emotion categories are distinct from more general dimensions of valence and arousal. We complement these analyses using a computational model of relational memory (the Tolman-Eichenbaum Machine^46^; TEM) that formulates how event sequences are represented in hippocampal-prefrontal systems. We train artificial agents to learn the structure of emotion-laden environments by binding sensory representations with structural knowledge about the world.

After training, we examine whether patterns of BOLD time-series in hippocampal-prefrontal systems covary with internal representations in computational agents, and whether they covary with human self-report. Together, these experiments test whether emotion concepts are represented in hippocampal-prefrontal systems, providing a neurocomputational explanation of how humans organize abstract emotion knowledge.

### Emotion categories and affective dimensions are represented in hippocampal-prefrontal systems

We first evaluated whether hippocampal activity represents individual emotion concepts in a format that is not merely based on variation in valence and arousal. To this end, we trained multivariate decoders that learn to map patterns of hippocampal fMRI time series (9,538 measurements in 1,816 voxels, per subject on average) onto multivariate emotion ratings. We fit separate models for emotion category and valence-arousal ratings using data acquired from multiple different film stimuli. Models were evaluated by testing them on independent stimuli (leave-one-film-out cross-validation, 13 films total for each of *N* = 29 subjects) and comparing the readout of category and valence-arousal decoders (see Methods). This analysis revealed that category ratings were more accurately decoded (*z =* 0.0525, 95% bootstrap *CI* [0.0454, 0.0592]) than valence-arousal ratings (*z =* 0.0286, 95% bootstrap *CI* [0.0197, 0.0389]; Δ*z =* 0.0239, 95% bootstrap *CI* [0.0183, 0.0297], *p* = .0002; Fig. 2b, left; see Supplementary Figs. 6 and 7 for contributions of individual voxels to the predictions), consistent with the proposal that hippocampal activity represents emotion concepts.

Because the hippocampus represents event sequences in a hierarchal manner^47–49^, sequences of emotional events should produce patterns of hippocampal activity in which emotions that tend to occur in temporal proximity (e.g., *disgust*, *anger*, and *guilt* or *anxiety*, *fear*, and *surprise*) exhibit more similar patterns of hippocampal response, so as to form a conceptual hierarchy^5,50,51^. We assessed whether this was the case by analyzing the outputs of hippocampal decoders using agglomerative clustering. For each subject, we computed the similarity of model predictions over time to produce a conceptual similarity matrix (Supplementary Fig. 2). We modeled the similarity structure in two ways: with a model that estimated temporal similarity by averaging across concepts of the same valence (referred to as the valence model), and with another model that estimated temporal similarity for all pairs of concepts (the full model; Fig. 1c). To estimate model generalizability, we performed a leave-one-subject-out procedure in which temporal correlations were averaged across all but one subject and compared to the similarity structure of the held-out subject. This generalization test revealed that the correspondence across participants in the fully specified model (Spearman’s *r* = .98, *SD* = .007, *p <* .0001) was greater than that of the valence only model (Spearman’s *r* = .84, *SD* = .013, *p <* .0001; Fig. 1d), and that a substantial portion of variance across participants remained after accounting for valence (Spearman’s *r* = .49, *SD* = .014, *p <* .0001). These results suggest that the hippocampus represents a hierarchy of emotion concepts that is stable across participants (Fig. 1b) and cannot be explained by high-level distinctions in valence alone.

**Figure 1.**
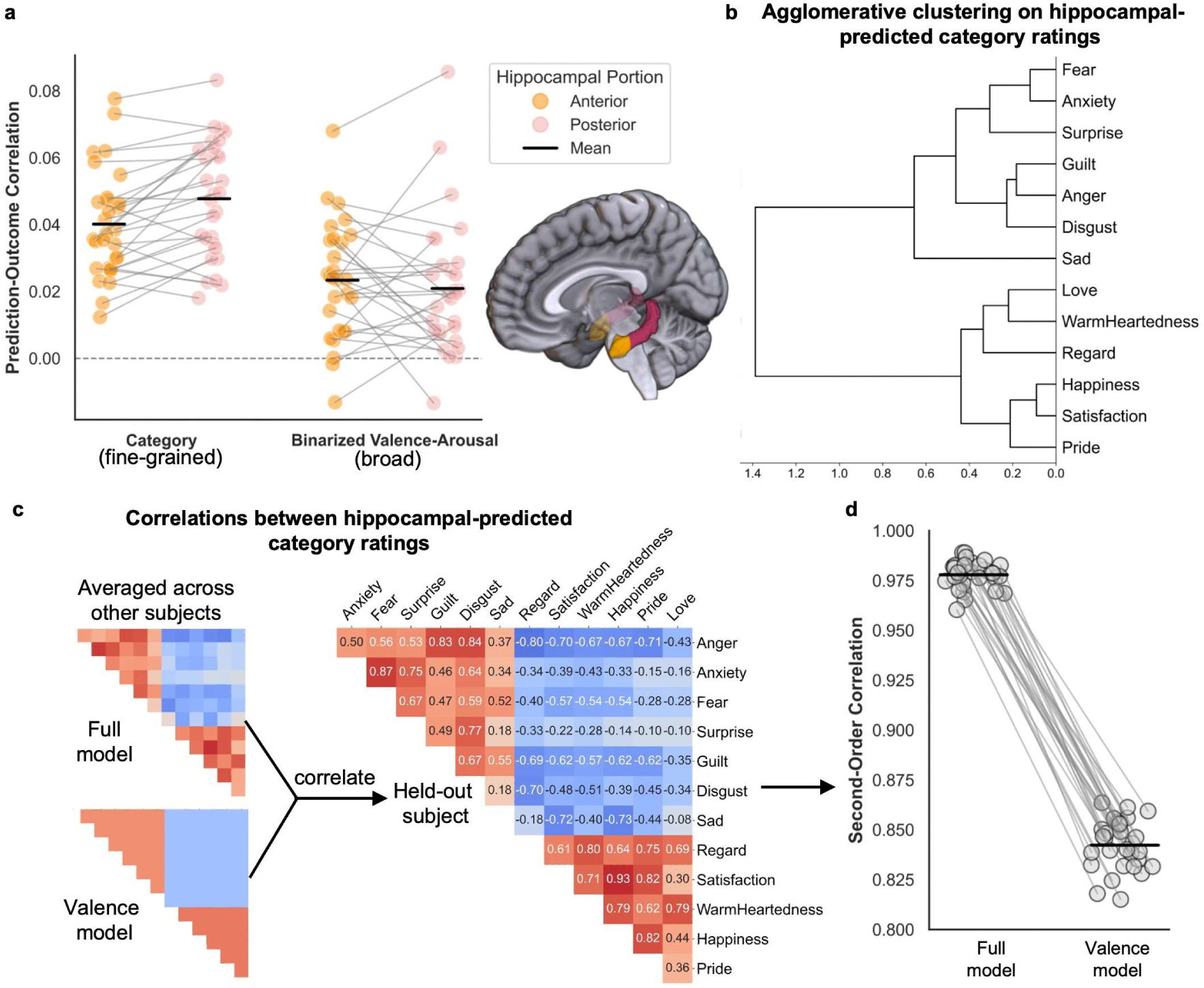
Decoding the representation of emotion concepts in the human hippocampus. **(a)** Performance of decoding models trained to predict emotion category and binarized valence-arousal ratings from BOLD activity in anterior and posterior hippocampus. Each point represents the prediction-outcome correlation averaged across rating items for category or binarized valence-arousal. Points from the same subjects are connected by gray lines, black horizontal lines indicate the mean, and gray dashed lines represent chance (i.e., zero correlation between predicted and observed outcomes). **(b)** Agglomerative clustering on group mean category ratings predicted from hippocampal BOLD activity. **(c)** Procedure for testing the structure of hippocampal representations across subjects. First-order correlation matrices illustrating the similarity structure of predicted emotion concepts from hippocampal fMRI patterns. Fully specified (all possible pairs of emotion concepts) and valence models (simplified models averaging within and between positive and negative categories) characterize covariation between emotions and are estimated using data from all but one hold-out subject (S01 shown here; see Supplementary Fig. 3 for all data). **(d)** Second-order correlations between each held-out subject’s similarity matrix and the full and valence models. Each point represents a subject. Points from the same subjects are connected by gray lines, black horizontal lines indicate the group mean.

Central to accounts of memory, navigation, and emotion processing^52^, the hippocampal long axis is characterized by gradients in connectivity, gene expression^53^, and behavioral specialization^54,55^. Particularly relevant for the representation of emotion knowledge are observations that hippocampal activity represents event sequences at multiple temporal scales, with more rapid changes coded in more posterior portions of the hippocampus^55^. Such variation leads to the prediction that fine-grained distinctions between emotional events that occur over shorter timescales should be captured by activity in posterior hippocampus, whereas more general distinctions that take place over longer timescales should be represented in more anterior portions of the hippocampus. To examine variation in emotion representation along the hippocampal long axis, we trained multivariate decoders to predict variation in fine-grained (e.g., *fear*, *anger*, *pride*, *joy* or *surprise)* and broad emotion concepts (e.g., *good,* or *activated*) using signals either in posterior or anterior hippocampus (based on a segmentation at the plane y = -21 in MNI space). Model comparisons revealed that decoding performance depended on both hippocampal portion and representational scale (Δ*z =* 0.0101, 95% bootstrap *CI* [0.0054, 0.0149], *p* = .0033, Fig. 1a), with greater decoding performance in posterior hippocampus as emotion granularity increased (see Supplementary Fig. 5 for performance on all ratings).

If the hippocampus conveys information about emotion concepts to entorhinal cortex and vmPFC to form a relational basis for emotion mapping, fMRI activity in these neocortical structures should more accurately track valence and arousal compared to signals in the hippocampus. Training and evaluating multivariate predictive models for these regions, we found that valence-arousal dimensions were more accurately decoded in vmPFC and entorhinal cortex compared to the hippocampus (Δ*z =* 0.0136, 95% bootstrap *CI* [0.0054, 0.0219], *p* = .0007). This between region difference was larger for decoding valence-arousal dimensions than emotion categories for entorhinal cortex (Δ*z =* 0.0130, 95% bootstrap *CI* [0.0067, 0.0193], *p* = .0008), and to a lesser extent vmPFC (Δ*z =* 0.0086, 95% bootstrap *CI* [0.0023, 0.0150], *p* = .0627; Fig. 2b). Thus, although all three regions contain information related to emotion concepts (see Supplementary Table 1 and Supplementary Figs. 6-9), neocortical areas contained relatively more information about affective dimensions than the hippocampus.

**Figure 2.**
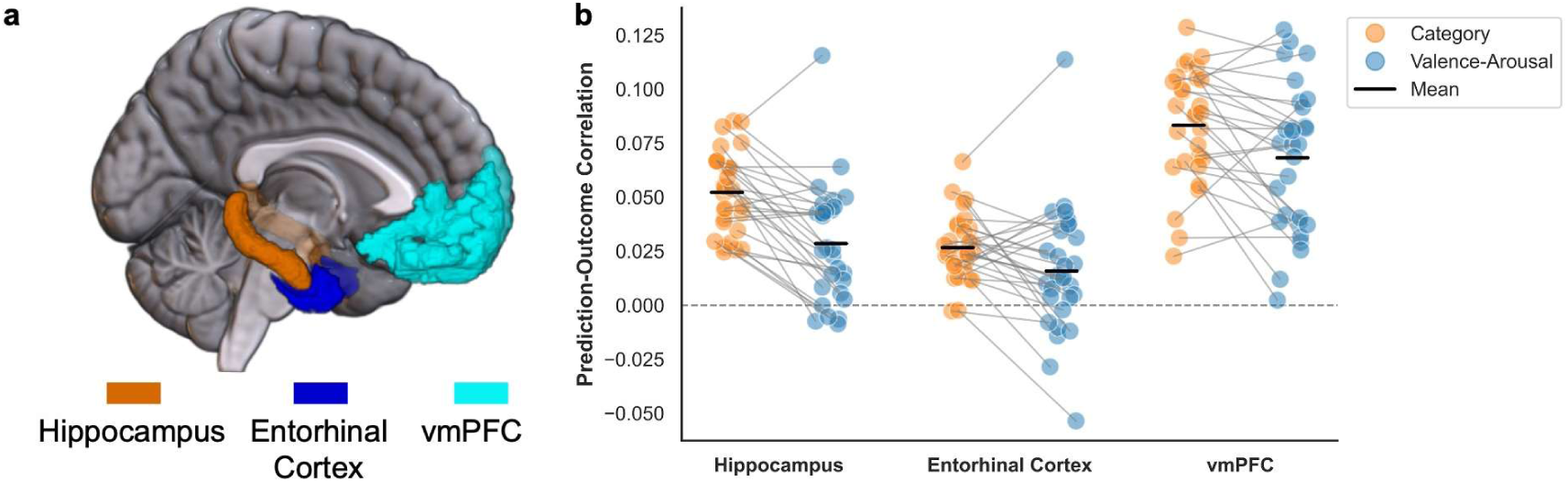
Decoding self-reported emotion in hippocampal-prefrontal systems. **(a)** Rendering of parcellations of hippocampus (orange), entorhinal cortex (dark blue), and ventromedial prefrontal cortex (vmPFC; light blue). **(b)** Performance of decoding models trained to predict emotion category and valence-arousal ratings from BOLD activity in hippocampus, entorhinal cortex, and vmPFC. Each point represents the prediction-outcome correlation averaged across rating items for category or valence-arousal. Points from the same subjects are connected by gray lines, black lines indicate the mean, and gray dashed lines represent chance.

### Hippocampal-prefrontal systems exhibit map-like representations of emotion concepts

The results of multivariate decoding experiments are consistent with the hypothesis that emotion concepts are represented in hippocampal-prefrontal systems. However, because predictive models do not reveal the specific format of the underlying representations (e.g., sparse representations of individual concepts or an integrated representation of valence and arousal as a two-dimensional space), it is possible that decoders leveraged signals other than the factorized representations of “what” and “where” hypothesized to underlie the formation of cognitive maps^56,57^. To probe the representational format of hippocampal activity more explicitly, we simulated concept formation using TEM, a computational model of relational memory (Fig. 3).

**Figure 3.**
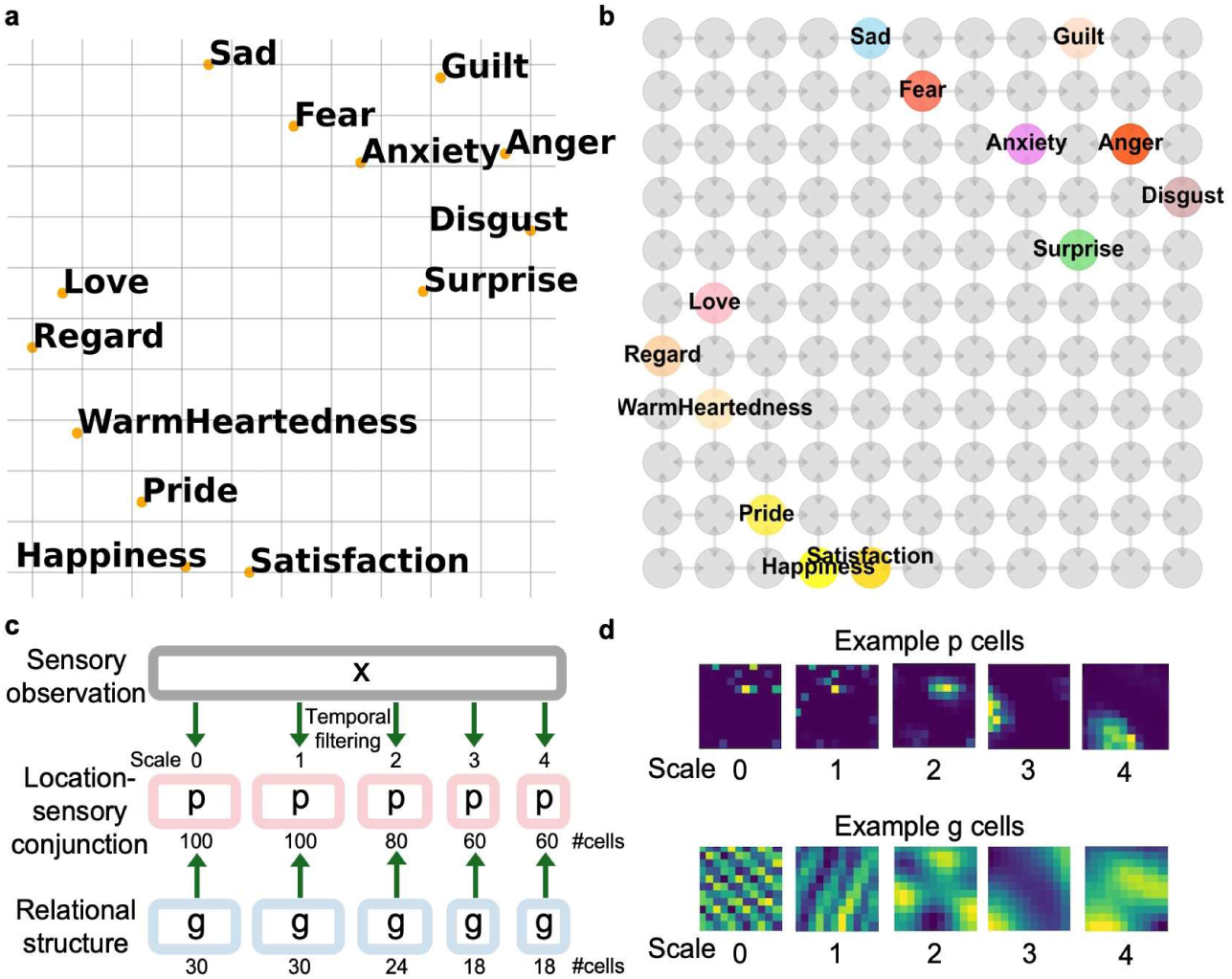
Simulating the construction of emotion maps using the Tolman-Eichenbaum Machine (TEM). **(a)** Multidimensional scaling of emotion category ratings across all film stimuli. Emotion categories are positioned relative to one another, with closer positions indicating higher correlations between ratings. **(b)** 11-by-11 discrete environment created for emotion concepts based on their locations in the multidimensional scaling of category ratings. Nodes with an emotion category assigned are colored and labelled. Arrows between nodes indicate possible movements of agents in the environment. **(c)** Schematic of layers and hierarchical organization of TEM. The model consists of two layers: **g**, which learns the relational structure between abstract locations in the environment, and **p**, which receives inputs from sensory observations and abstract locations to learn their associations. Different temporal filters for sensory observations are learned by separate streams of the model, capturing varying scales of the spatiotemporal structure. **(d)** Rate maps of example hippocampal and entorhinal cells obtained by averaging the activity of each cell at each node after training.

In learning to map the environment, TEM factorizes sensory observations into two separate bases—one that represents the contents of experience and another that represents relationships between experiences—and binds them together into a conjunctive code. This conjunction is represented in layer **p**, which is used to make predictions about upcoming sensory observations and locations in layer **g**. Whereas the activity in layer **p** resembles hippocampal population activity that encodes “what” is located “where” in an environment, activity in layer **g** resembles that of entorhinal cortex, representing an abstract structural code that can be used for making inferences in any environment or in new problems that have a similar relational structure.

We created an environment based on the covariance of emotion categories derived from film ratings so that agents exploring the environment would experience transitions between emotional events that approximate human experiences. We used multidimensional scaling to map emotion categories into a two-dimensional affective space, which was discretized into a grid environment. During training, agents randomly explored this environment, learning the relational structure of emotion concepts through experience, mimicking one way in which the brain could learn to represent emotion concepts over time. This simulation enabled us to define representations of emotion concepts and relations between them and evaluate how accurately they can be decoded from patterns of hippocampal activity in the fMRI signal. To model human brain responses during film viewing, we generated TEM activation time series by averaging the activation of layers **p** and **g** in artificial agents as they navigated the affective space, weighting the responses in layers **p** and **g** using a linear combination of ratings on all emotions at each point.

Similar to the multi-scale representations observed in place cells across the hippocampal long axis^58,59^, TEM includes artificial neurons at multiple levels of abstraction, allowing it to represent both fine-grained and coarse aspects of the environment. Because representations in layers **p** and **g** differed the most at small scales in our simulations (Fig. 3d, and Supplementary Fig. 10), we predicted that hippocampal decoding of layer **p** would be more accurate than layer **g** for units with narrow firing fields (smaller scales), as they better capture transitions between specific emotions as opposed to large transitions in the affective space (larger scales). Consistent with this prediction, we found higher cross-validated decoding performance for layer **p** than **g** (Δ*z =* 0.0017, 95% bootstrap *CI* [0.0001, 0.0034], *p* = 0.0145; Fig. 4a). Further, a linear contrast across representational scales revealed that as scale decreased, decoding performance increased more for layer **p** than for layer **g** (Δ*z* = 0.0352, 95% bootstrap *CI* [0.0254, 0.0448], *p* < .0001; Fig. 4c, left), indicating that more accurate decoding of layer **p** activity was driven by information related to small-scale conjunctive codes.

**Figure 4.**
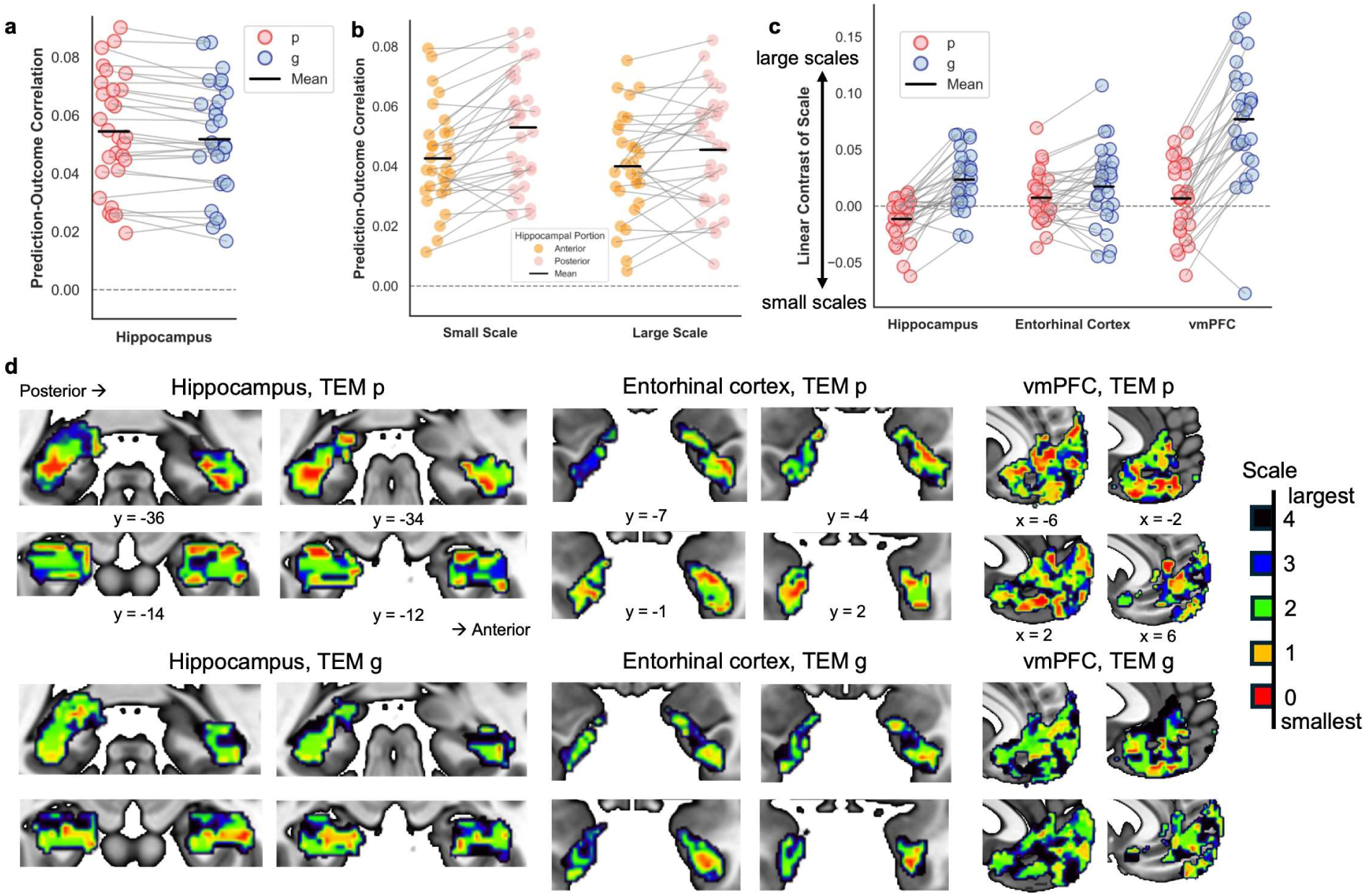
Decoding representations of conjunctive codes and relational structure in hippocampal-prefrontal systems. **(a)** Performance of decoding models trained to predict activity in layers **p** and **g** of the Tolman-Eichenbaum Machine (TEM) from BOLD activity in the hippocampus. Each point represents the prediction-outcome correlation averaged across cells and scales for one subject. **(b)** Performance of decoding models trained to predict small- and large-scale **p** activity from BOLD signal in anterior and posterior hippocampus. Each point represents the prediction-outcome correlation for one subject. **(c)** Linear contrast of decoding performance across TEM scales. Negative values indicate higher decoding performance at smaller scales; positive values indicate higher performance at larger scales. See Supplementary Fig. 11 for decoding performance at each individual scale. **(d)** Maps indicating the scale with the largest PLS coefficient, with smaller scales shown with warm colors and large scales shown with cool colors (see also Supplementary Fig. 12). Maps indicate the scale with the largest *t*-statistic (FDR *q* < 0.05) averaged across cells for each layer (Supplementary Figs. 13-15). In the hippocampus, more warm colors are shown in the posterior portions for **p** relative to **g**. In the ventromedial prefrontal cortex (vmPFC), more cool colors are shown for **g** relative to **p**.

Because firing field size varies along the hippocampal long axis^58,59^, we next tested whether information about smaller-scale layer **p** activity was more strongly represented in posterior hippocampus. To this end, we trained separate decoding models to predict TEM activations at small (0 and 1) and large representational scales (2 through 4), and compared the accuracy of readouts from anterior versus posterior hippocampal fMRI activity. This analysis revealed differences in decoding accuracy that depended on both hippocampal portion and scale (Δ*z =* 0.0052, 95% bootstrap *CI* [0.0033, 0.0071], *p* = .0019; Fig. 4b and 4d, left), indicating that smaller-scale layer **p** activity was decoded more accurately in posterior than anterior hippocampus.

We observed that hippocampal decoders predicted variation in emotional experience as reflected in both human self-report and the activity of TEM agents. To determine whether decoders utilized common patterns of hippocampal response, we compared decoding performance before and after controlling for predictions of **p** (see Methods, ‘Comparisons of decoding performance of TEM activity’ for details). We found that accounting for hippocampal signals related to layer **p** activity impaired the readout of emotion category ratings (Δ*z*_cat_ *=* 0.0040, 95% bootstrap *CI* [0.0020, 0.0061]) more so than valence-arousal ratings (Δ*z*_dim_ *=* 0.0002, 95% bootstrap *CI* [0.0010, 0.0014]; Δ*z*_cat-dim_ *=* 0.0039, 95% bootstrap *CI* [0.0021, 0.0057], *p* = .0010). This result suggests that a conjunctive representation of emotion concepts can account for some, but not all, of enhanced prediction of emotion category ratings compared to valence-arousal dimensions in the hippocampus.

The entorhinal cortex and vmPFC are thought to represent relations among objects, locations, and goal-states in a domain general way^36,39,60,61^. If it plays a similar role in representing relations among emotion concepts, we would expect that decoding layer **g** activity in these regions should be more accurate than decoding layer **p** activity. We tested this prediction by comparing the decoding accuracy as a function of brain region (hippocampus, entorhinal cortex, and vmPFC), layer (**p** and **g**), and scale (small and large). Analysis of variance revealed that decoding performance depended on all three variables (a three-way interaction: *F*(8, 616) = 12.44, *p* < .0001; Supplementary Fig. 11). Direct comparisons indicated that there was better readout of layer **g** than layer **p** activity at large scales in vmPFC (Δ*z =* 0.0712, 95% bootstrap *CI* [0.0614, 0.0809], *p* < .0001; Fig. 4c, right), and to a lesser degree in the entorhinal cortex (Δ*z =* 0.0099, 95% bootstrap *CI* [0.0001, 0.0199], *p* = .0461; Fig. 4c, middle). Decoding layer **g** was more accurate at larger scales in the vmPFC than in both hippocampus (Δ*z =* 0.0542, 95% bootstrap *CI* [0.0447, 0.0640], *p* < .0001; Fig. 4c, left and right) and entorhinal cortex (Δ*z =* 0.0604, 95% bootstrap *CI* [0.0506, 0.0700], *p <* .0001; Fig. 4c, middle and right). Conversely, decoding layer **p** activity was better at small scales in the hippocampus compared to both entorhinal cortex (Δ*z =* 0.0191, 95% bootstrap *CI* [0.0093, 0.0289], *p* = .0003; Fig. 4c, left and middle) and vmPFC (Δ*z =* 0.0182, 95% bootstrap *CI* [0.0086, 0.0279], *p* = .0031; Fig. 4c, left and right). Together, these results suggest that the vmPFC contains more information about large-scale structural abstractions (e.g., that one portion of a video was more pleasant than another), whereas the hippocampus represents more information about small-scale representations of individual concepts.

## Discussion

Knowledge about the world is thought to be organized into a map-like representation that enables organisms to flexibly navigate complex environments^62^. In these accounts, domain general cortico-hippocampal networks use similar mechanisms to organize knowledge of physical locations^31^, sensory percepts^36^, social relationships^37^, and abstract concepts^39^. We found that patterns of human hippocampal activity represent multiple emotion concepts hierarchically in a way that was not reducible to more basic dimensions of valence and arousal, suggesting that emotion knowledge may be organized via similar mechanisms as other domains.

Several prominent accounts of emotional experience suggest that the hippocampus and vmPFC, as nodes in the default network, are involved in the conceptualization of emotional experiences^63,64^. Consistent with these accounts and past decoding studies^17,18,22–24^, we found that information about emotion categories and valence-arousal dimensions was represented in multiple regions of the default network. Our observation that hippocampal representations generalized across film stimuli with different audiovisual and linguistic features suggests that emotion concepts may result from the abstraction of sensory signals in the environment that are shared across individuals. Further supporting this view, the engagement emotion concepts we decoded were based on self-report ratings from independent observers. Nevertheless, whether emotion concepts represented in hippocampal-prefrontal systems reflect the statistical regularities of important life events that are shared across individuals and cultures, or whether they are constructed more flexibly as *ad hoc* categories^65^ shaped by individual experience, remains an open question. Existing developmental research^24^ suggests that neural representations of emotion concepts are present early in life, as young as age five, and that they stabilize during adolescence, consistent with a learning process that extracts regularities over time. However, longitudinal research is needed to determine the extent to which behaviors that draw on emotion knowledge follow a similar developmental trajectory as the hippocampus.

Our findings go beyond results of existing decoding studies by demonstrating that an unsupervised agent using conjunctive coding—which binds sensory information and structural knowledge about emotions (i.e., that some experiences feel more or less pleasant or arousing than others)—can model hippocampal-prefrontal responses to emotional films that are associated with self-reported emotion. Whereas past work has shown that information about different aspects of emotional experiences is distributed across the cortex^22,23^, and that differences in valence covary with BOLD responses in cortical midline regions^17,18^, our computational experiments advance an account of how the human brain organizes emotion concepts. We found that hippocampal signals represented locations in affective space produced through a conjunctive process (as reflect in layer **p** activity in TEM), whereas vmPFC signals were most strongly related to large scale bases that span affective space (i.e., grid-like layer **g** activity with low frequency firing fields, see Fig 3d). More accurate readout of large-scale layer **g** activity in vmPFC aligns with its longer temporal integration windows, a feature typical of brain regions at the top of the cortical hierarchy^66,67^. Considered alongside evidence that vmPFC and other nodes of the default network exhibit grid-like activity during tasks from multiple domains^36,37,68^, these findings suggest that similar neural mechanisms may be involved for reward-based decision making^69^, social inferences^70^, and reporting emotional experience^18^–all of which require relational processing.

We observed variation in the granularity of emotion concepts along the hippocampal long axis. This finding aligns with observations that the posterior hippocampus encodes fine-grained distinctions between entities^54^, characterized by smaller, more precise firing fields compared to anterior hippocampus^58,59^. It is also consistent with evidence that the hippocampus represents event sequences hierarchically—with longer timescales being represented in more anterior portions of the hippocampus^55^. Notably, post hoc analyses revealed that several emotion categories (e.g., *love*, *satisfaction*, *guilt*) were better decoded from anterior compared to posterior hippocampus (Supplementary Fig. 5 and Supplementary Table 2). This pattern could reflect the temporal dynamics of these emotions, which often involve the integration of various aspects of emotional events over longer timescales than putatively basic emotions which are thought to occur with a rapid onset and short duration^71^.

Unexpectedly, we found more signal related to emotion categories than valence-arousal dimensions across hippocampus, entorhinal cortex, and vmPFC. Several factors might explain why we did not observe stronger representation of dimensional variables in entorhinal cortex and vmPFC as expected. First, the naturalistic film stimuli lacked sharp transitions between emotion categories that were independent of valence, which may have limited our ability to separately manipulate these variables. Future research using more structured emotional stimuli (e.g., controlled trajectories of transitions in the valence-arousal space) could reveal clearer functional dissociations between regions. Even so, one would expect the information content to be somewhat similar across hippocampal-prefrontal systems regions they reciprocally convey information^72^ about emotion concepts and their relations to organize conceptual knowledge.

Using computational models to characterize neural responses to naturalistic stimuli provides a fresh look on how the brain constructs cognitive maps more generally. Previous investigations have relied on synthetic stimuli presented in an impoverished sensory environment^36,39^; our use of naturalistic movie stimuli offers a more ecologically valid window into how hippocampal-prefrontal systems process dynamic, rich, emotional experiences. Further, our application of TEM^46^ to human fMRI data bridges theoretical models, animal electrophysiology, and human neuroimaging. This approach allows us to make inferences about potential computational mechanisms underlying the observed representational patterns, connecting macroscale BOLD signals to principles consistent with those learned from cellular-level organization. However, TEM is not the only model of hippocampal computations. Alternative models grounded in reinforcement learning^73^ may offer complementary insights, particularly for studying emotion concepts, which inherently encode value and motivational relevance. This framework aligns with evidence suggesting the role of the anterior hippocampus and vmPFC in motivation and goal-directed behavior^74–76^. Future research should examine whether models of reinforcement learning can better predict hippocampal-prefrontal representations of emotion concepts.

Our findings extend our understanding of hippocampal concept cells, which have primarily been linked to physical entities (e.g., persons, objects) in single-unit recording studies^41,77^. Whereas prior fMRI studies have explored hippocampal-prefrontal contributions to the processing of abstract information, they have focused largely on grid-like coding patterns when participants traversed conceptual spaces^36,37^. Here, we reveal that the hippocampus represents emotion concepts at multiple levels of abstraction, dovetailing with recent findings demonstrating that its representational capacity spans both physical and abstract domains.

Importantly, our decoding models generalize across different films, demonstrating that these representations are not tied to low-level perceptual features but instead capture higher-level, abstract properties of emotion concepts. This generalizability highlights the hippocampus’ role in encoding conceptual structure beyond specific sensory experiences, further supporting its role in the organization of abstract knowledge. In sum, the current work demonstrated that hippocampal-prefrontal systems exhibit map-like representations of emotion concepts. Our findings offer new insights into the long-standing observation that people report their feelings using a mental map organized by dimensions of valence and arousal^1–4^. Although it has long been suggested that our conceptualization of emotions emerges from these fundamental dimensions^65,78^, we have shown that brain systems responsible for computing relations between distinct experiences are capable of structuring emotion knowledge in a low-dimensional affective space. These findings raise the possibility that the map-like structure observed in self-report is the product of computations performed in hippocampal systems, rather than being an innate structure afforded by the human brain.

## Methods

### Emo-FilM dataset

Both the fMRI data (https://openneuro.org/datasets/ds004892/versions/1.0.0) and emotion ratings (https://openneuro.org/datasets/ds004872/versions/1.0.1) used in this work are from the Emo-FilM dataset^45^ publicly available on OpenNeuro. Both studies were approved by the Geneva Cantonal Commission for Ethics in Research.

#### fMRI data

Brain activity was measured using fMRI in 30 healthy subjects that watched 14 short films (mean duration 11 min 26 s) in a pseudo-random order over four sessions (see Morgenroth et al., 2024^45^ for detailed descriptions of the films). All subjects were right-handed and met the inclusion criteria of normal or corrected-to-normal vision, high-level English comprehension, no history of psychiatric or neurological illness, and no neuropharmacological or recreational drugs. We excluded subject 07 due to missing data from 3 films on OpenNeuro, leaving 29 subjects (18 females, ages 18-34). MRI data were acquired on a 3T Siemens Magnetom TIM Trio scanner with a 32-channel head coil (Siemens, Erlangen, Germany). Anatomical T1 images were acquired with a GRAPPA sequence for the purpose of co-registration (TR = 1.9 s, TE = 2.27 ms, flip angle = 9°, FOV = 256 mm, resolution = 1 mm^3^, in plane resolution of 256 × 256 × 192 sagittal slices). Functional images were acquired with the same multi-band frequency protocol (TR = 1.3 s, TE = 30 ms, flip angle = 64°, FOV = 210 mm^2^, resolution = 2.5 mm^3^, 54 interleaved slices).

#### fMRI preprocessing

Preprocessing of fMRI data, performed using FEAT (FMRI Expert Analysis Tool) from FSL, included co-registration to structural, standard space, and each subject’s functional volume, motion correction, non-brain tissue removal, spatial smoothing (6 mm FWHM), grand-mean intensity normalization, high-pass temporal filtering (50 s cutoff), and regression of white matter, cerebrospinal fluid, and six motion parameters.

#### Emotion ratings

Emotion ratings were collected from 44 independent participants (23 females, aged 20-39) recruited online, using the same inclusion criteria as the fMRI sample. Participants were from the Geneva area, matched with the fMRI sample, but with no overlap between the two samples. A total of 55 items were rated across six groups of items: Appraisal, Expression, Physiology, Motivation, Feeling, and Discrete Emotion. Participants completed the rating tasks at their own pace within a six-week period, using their own computers. To rate each item, participants continuously moved a mouse-controlled cursor along a bar (sampling rate = 1 Hz). Each participant rated six items in total, with one item rated across all films before moving to the next item, ensuring a sufficient delay between repetitions of the same film. Consensus ratings were derived by averaging from three or four raters per item and film (see Morgenroth et al., 2024^45^ for a detailed description of consensus calculation and quality control procedures).

In the current study, we used all 13 items (*anger*, *anxiety*, *fear*, *surprise*, *guilt*, *disgust*, *sad*, *regard*, *satisfaction*, *warmheartedness*, *happiness*, *pride*, *love*) in the Discrete Emotion category. For valence and arousal, we selected four items (valence: *good*, *bad*; arousal: *calm* (*restless*-*calm*), *at ease* (*nervous*-*at ease*) from the Feeling items, as they best align with our goal of capturing emotion concepts in terms of their location within the valence-arousal space. Other Feeling items including *intense emotion*, *strong* (*weak*-*strong*), and *alert* (*tired*-*alert*) were not used because they were less directly relevant to arousal or did not align with the goal of capturing conceptual knowledge of emotion rather than physiological states. For example, *intense emotion* reflects the magnitude of emotions, *weak*-*strong* may reflect a sense of capability and self-efficacy, and *tired*-*alert* is more indicative of wakefulness rather than the concept of arousal.

### Defining regions of interest

Masks of the hippocampus and entorhinal cortex were obtained from the Julich-Brain Cytoarchitectonic Atlas^79^. Masks of anterior and posterior hippocampus were obtained by splitting the hippocampal mask at y = −21 in the MNI space^80^. For the vmPFC mask, we selected Brodmann areas 10, 11, 14, 24, 25, and 32^17^, obtained from a multimodal parcellation of the human cortex.

### Simulating hippocampal learning with the Tolman-Eichenbaum Machine

TEM is an artificial neural network trained to navigate on a connected graph^46^. At each time step, it takes the current sensory observation (**x**) and an action (**a**) as inputs, and outputs a predicted sensory observation for the next time step. To accomplish the objective of accurately predicting next sensory inputs even when taking paths that have not been experienced before, TEM factorizes its representation into two parts: 1) relations between locations on the graph and 2) the association between sensory observations and their location on the graph.

TEM factorizes sensory information using two layers that are inspired by the functional anatomy of the entorhinal cortex and hippocampus. The entorhinal layer of the network (**g**) generates distinct representations for each location and learns their relations through path integration. At each time step, **g** updates its representation based on the received action, using a recurrent architecture with weights updated via backpropagation during training. The hippocampal layer of the network (**p**) receives sensory input along with the estimated location from **g**. Memories of sensory-location associations is stored in Hebbian weights between units in **p** and can later be retrieved by attractor dynamics.

To efficiently represent environments, TEM is organized into multiple parallel streams, each capturing information at different temporal and spatial scales. Each stream processes input separately, applying a unique temporal smoothing filter, and transitions via path integration (see Whittington et al., 2020^46^ for details). These streams remain distinct during learning but integrate during memory retrieval through attractor dynamics. Lower-index streams encode finer-grained details, while higher-index streams capture broader structure.

To enable TEM to learn a relational structure that generalizes across different environments, we trained the model in 16 distinct 11×11 grid environments, each with randomly generated 45-dimensional one-hot sensory inputs. At each time step, the agent could take one of five possible actions—moving up, down, left, right, or staying still—with equal probabilities. The results presented in the main text are based on the model trained for 32,000 iterations, at which point the loss and performance in predicting next sensory observations began to plateau (Supplementary Fig. 16). Additional analyses using models trained for 42,000 and 50,000 iterations showed consistent results (Supplementary Fig. 17 and Supplementary Tables 3-6), confirming the robustness of our findings.

### Creating the TEM environment for emotion concepts

We first calculated the Pearson correlations of time series of ratings between each pair of emotion categories concatenated across all 14 films. Next, we computed the dissimilarity between pairs of emotion categories using 1 − Pearson correlation and performed metric multidimensional scaling^81^ (MDS) on the dissimilarity matrix. The resulting two-dimensional MDS space was discretized into 121 locations, forming an 11 × 11 grid, to match with the environments used during training. Each emotion category was assigned to the location closest to its MDS coordinate. To avoid overlap, if a location was already occupied by another category, the current category was assigned to the nearest available, unoccupied location.

To ensure that the MDS solution accurately reflected the (dis)similarities between emotion categories as indicated by the ratings, we repeated the process 500 times using different random seeds for the MDS procedure. We selected the seed that resulted in the lowest Spearman correlation between the city-block distances between category pairs in the environment (which correspond to the minimum number of time steps needed to transition from one category to another) and the Pearson correlations derived from the ratings.

As in the training environment, the agent in the emotion environment could take one of five actions—up, down, left, right, or staying in the same location—each with equal probability. To ensure that each emotion category had a unique sensory input, we first randomly assigned a distinct 45-dimensional one-hot code to each emotion category. For locations without an associated emotion category, we assigned random 45-dimensional one-hot codes that did not overlap with those of the emotion categories.

### Simulating emotional experience with TEM

With trained TEM model weights, the agent walked randomly in the environment created for emotion concepts for 12,100 steps (to roughly sample each of 121 locations 100 times). For each emotion category, we calculated the average activations of all units in **p** and **g** across the second half of the steps to ensure that the agent had enough experience to develop a stable representation of the new environment it had never encountered during training. To generate a time series of responses to emotion concepts throughout the films, we computed the weighted average activation across all categories, as we assume that multiple emotion concepts can be coactivated during film viewing. Specifically, we weighted the activations of units in layers **p** and **g** based on emotion ratings at each time point, such that segments rated more highly on certain emotions resulted in stronger activations for those concepts, while minimizing contributions from unrelated emotions.

### Multivariate decoding of BOLD fMRI

We specified partial least squares (PLS) regression models (SIMPLS algorithm^82^) with 20 components separately for decoding patterns of BOLD activity to predict emotion category ratings, valence-arousal ratings, and activation in different layers of TEM (Supplementary Fig. 1). Fitting separate models for different outcome variables simplifies the interpretation of model performance, because the different outcome blocks (e.g., category ratings and valence-arousal ratings) are correlated and could influence the estimation of betas for different outcome variables^83,84^. For all models, the predictor block consisted of an *n*_timepoint_ × *p*_voxel_ dimensional matrix of BOLD signal (*n* = 9,538 timepoints; *p_hippocampus_*= 1,816 voxels, *p_entorhinal cortex_* = 711 voxels, *p_vmPFC_* = 5,597 voxels). Outcome blocks consisted of *n*_timepoint_ × *p*_item_ matrix of ratings (*p* = 13 category ratings or *p* = 4 dimensional ratings), or a *n*_timepoint_ × *p*_unit_ matrix of TEM activity (*p****_p_*** = 400 units, *p****_g_*** = 120 units). To align with BOLD sampling, time series of the outcomes (ratings and TEM activity) were resampled to match the sampling rate of BOLD data (i.e., 1/1.9 Hz) and convolved with a canonical double gamma response function to account for hemodynamic delay^85^. For each of the three regions of interest, we therefore developed four models (category, valence-arousal, **p**, and **g**). To test the granularity of emotion concepts and the scale of **p** represented along the hippocampal long axis, we additionally specified separate PLS models separately decoding signals from the anterior and posterior hippocampus (*p_anterior hippocampus_* = 745 voxels, *p_posterior hippocampus_* = 1,139 voxels) to predict emotion category, binarized valence-arousal, small-scale **p** (*p****_p_*** = 200 units), and large-scale **p** (*p****_p_*** = 200 units).

We quantified decoding performance using leave-one-film-out cross-validation in which PLS regression models were trained on 13 films and tested on the left-out film. The predicted outcome time series in the test fold were correlated with the observed time series of the outcomes, and performance was calculated as the average correlation across the 14 validation sets (i.e., films) for each subject. To test whether self-report ratings predicted from BOLD activity were partially explained by the decoded TEM activity, we used the time series of decoded TEM activity to predict the ratings using PLS. We then performed a partial correlation analysis between the decoded ratings and the actual ratings, while controlling for the ratings predicted by the decoded TEM activity.

### Characterizing the representational geometry of emotion concepts in the hippocampus

To assess the hierarchical organization of emotion concepts in hippocampal representations, we computed the similarity between predicted rating time series for each pair of emotion categories using Pearson correlation coefficient. We then applied agglomerative hierarchical clustering using the average linkage method to the dissimilarity matrix (1 – Pearson correlation). This procedure was repeated for each subject, and the resulting dendrograms were used to visualize the hierarchical relationships among emotion concepts.

To examine whether a shared conceptual similarity structure was present across subjects and whether it could be explained by valence, we implemented a leave-one-subject-out analysis. For each iteration, we averaged the pairwise category correlations across all subjects except the one held out, creating a group-level model of similarity between emotion concepts. Two group-level models were defined: a full model consisting of the mean correlation for each pair of the 13 categories, and a valence model, in which category pairs were grouped by valence (positive– positive, negative–negative, and positive–negative) and averaged within each group. We assessed model fit by computing Spearman rank correlations between the held-out subject’s pairwise category correlation matrix and each group-level model. To test whether the full model explained variance in the subject’s similarity structure beyond what could be attributed to valence, we regressed out the ranked values of the valence model from the subject’s category correlations and then computed a Spearman correlation between the residuals and the full model. Inference on group-level correlation coefficients was performed using t-tests against zero.

### Comparing multivariate decoding performance across subjects

We specified multiple linear mixed-effects models to compare decoding performance across models trained for different objectives (e.g., for categorical and dimensional self-report items) and using data from different regions (e.g., hippocampal and entorhinal regions of interest).

#### Comparing the decoding performance of self-report ratings

We specified a linear mixed-effects model to analyze decoding performance of self-reported ratings (i.e., Fisher z-transformed prediction-outcome correlation) as a function of rating item (including both category and valence-arousal), region, and their interaction as fixed effects. Random intercepts were included for subjects nested within rating items and regions. To compare decoding performance of category versus valence-arousal in the hippocampus, we contrasted the estimated marginal means of the hippocampus, by taking the average of 13 category items and subtracting the average of the four valence-arousal items, while setting other regions to zero. To compare how the decoding performance of valence-arousal in the hippocampus differed from that in the entorhinal cortex and vmPFC, we contrasted the average estimated marginal means of the four valence-arousal items in entorhinal cortex and vmPFC against that of the hippocampus (entorhinal cortex + vmPFC – hippocampus) for valence-arousal items only, while setting category items to zero. To compare how the difference in rating type (emotion category vs. valence-arousal dimension) varied across brain regions, we tested the interaction between rating type and region (hippocampus - entorhinal cortex or hippocampus - vmPFC).

To compare decoding performance between emotion category and binarized valence-arousal items in the anterior and posterior hippocampus, we specified a linear mixed-effects model with rating item, hippocampal portion, and their interaction as fixed effects, and random intercepts for subjects nested within rating items and hippocampal portions. We contrasted the estimated marginal means for emotion type and hippocampal portion. For the factor emotion, we subtracted the average of the four binarized valence-arousal items from the average of the 13 category items. For the factor hippocampal portion, we subtracted performance in the anterior hippocampus from that of the posterior hippocampus. We tested the interaction between these two factors to determine whether the difference in hippocampal portions (posterior - anterior) varied across emotion types/granularities (category - binarized valence-arousal).

#### Comparing the decoding performance of TEM activity

We specified a linear mixed-effects model to analyze decoding performance of TEM activity as a function of brain region, TEM layer, scale, and their interactions as fixed effects. Random intercepts were included for subjects nested within regions, TEM layers and scales. In these models, we averaged decoding performance across units for each combination of brain region, TEM layer and scale with subjects. This approach simplified the models and ensured a more appropriate estimation of degrees of freedom.

To compare the decoding performance of the **p** and **g** layers of TEM in the hippocampus, we contrasted the estimated marginal means of the hippocampus, by taking the average of all the scales of **p** and subtracting the average of all the scales of **g**, while setting other regions to zero. To examine trends in decoding performance across different scales (0 through 4), we specified a linear contrast on the estimated marginal means, assigning weights from −2 to 2 to represent the change in performance from small to large scales. This contrast was performed separately for each combination of brain region and TEM layer. We then conducted pairwise comparisons to assess the differences in the estimated marginal means of the linear contrast on scale between each pair of regions separately for each TEM layer and between each pair of TEM layers separately for each region.

To compare the performance of models trained to predict the activity of small- and large-scale units in layer **p** of TEM from signals in the anterior and posterior hippocampus, we specified a linear mixed-effects model with representational scale, hippocampal portion, and their interaction as fixed effects, with random intercepts for subjects nested within scale and hippocampal portion. We contrasted the estimated marginal means for the factors scale and hippocampal portion. For the factor scale, we subtracted the average of the three large scales (i.e., 2, 3, 4) from the average of the two small scales (i.e., 0, 1). For the factor hippocampal portion, we subtracted decoding performance in the anterior hippocampus from that of the posterior hippocampus. We tested the interaction between these two factors to determine whether the difference in decoding performance between hippocampal portions (posterior - anterior) varied across scales (small - large).

To compare the correlation between BOLD-predicted and observed self-report ratings after controlling for TEM **p** in the hippocampus, we specified a linear mixed-effects model with rating item, correlation type (correlation vs. partial correlation controlling for TEM **p**), and their interaction as fixed effects, with random intercepts for subjects nested within rating items and correlation types. We contrasted the estimated marginal means for the factors correlation type and hippocampal portion. For the factor correlation type, we subtracted the average of the partial correlations from the average of the correlations. For the factor emotion, we subtracted the average of the four valence-arousal items from the average of the 13 category items. We tested the interaction between these two factors to determine whether the difference in correlation type (correlation - partial correlation) varied across emotion types (category - valence-arousal).

We assessed the statistical significance of all effects of interest using a nonparametric permutation test based on sign flipping^86^. Specifically, for each permutation, we randomly flipped the sign of all data within each subject, thereby preserving the within-subject correlation structure while assuming exchangeability of subjects under the null hypothesis. This procedure maintains the dependency across repeated measures for each subject (e.g., brain regions, emotion ratings, TEM layers, TEM scales) and permits inference at the group-level. For each randomized dataset, we refit the linear mixed-effects model and extracted estimates of interest. The randomization null distribution was constructed from 10,000 permutations, and two-tailed *p*- values were computed as the proportion of permuted estimates that were more extreme than the observed estimate. To additionally provide robust uncertainty estimates, we computed 95% confidence intervals using 10,000 parametric bootstrap simulations^87^. These simulations generated synthetic observations based on parameter estimates from the original model. Mixed effects models were fit for each bootstrap iteration, producing a distribution of estimates use to derive 95% confidence intervals using the percentile method^88^.

## Author Contributions

P.A.K. and Y.M. contributed to designing the research, analyzing the data, and writing the paper.

## Data Availability

The fMRI data are available at https://openneuro.org/datasets/ds004892/versions/1.0.0. The emotion rating data are available at https://openneuro.org/datasets/ds004872/versions/1.0.1.

## Code availability

Code for all analyses will be made available upon publication at https://github.com/ecco-laboratory/EmotionConceptRepresentation. Code for training TEM model is adapted from https://github.com/jbakermans/torch_tem. The MATLAB interface used for creating TEM environments is available at https://github.com/jbakermans/WorldBuilder.

