## Supplementary for "The representation of emotion knowledge in hippocampal-prefrontal systems"

### Supplementary Figures

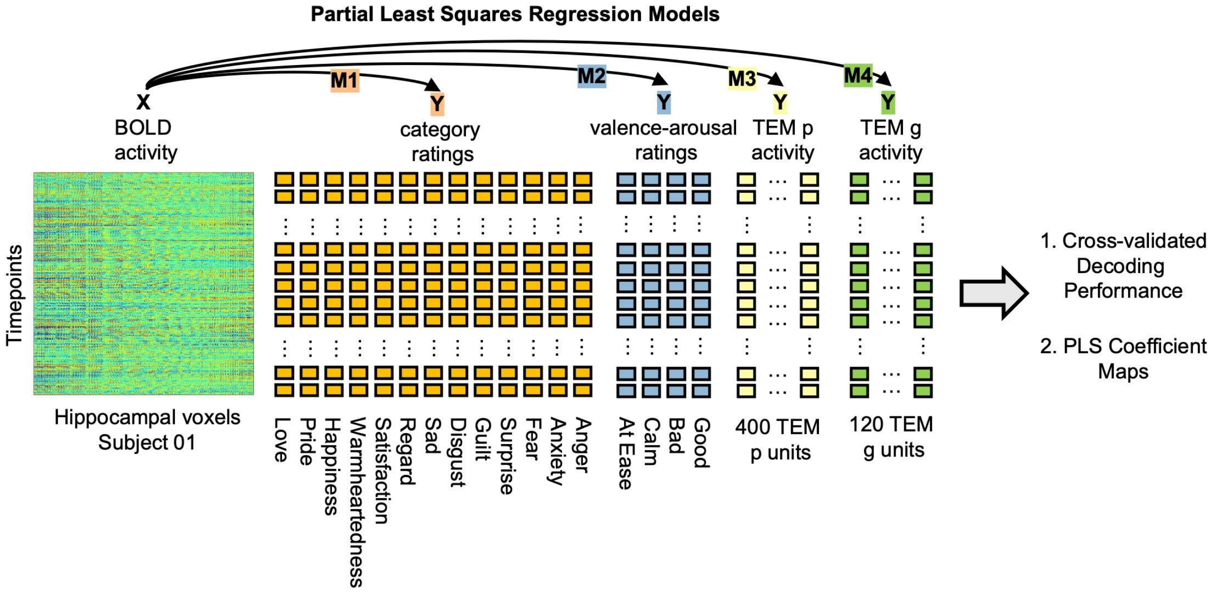


**Supplementary Figure 1. Schematic multivariate brain decoding.** Separate partial least squares (PLS) regression models were used to decode category ratings (shown in orange), valence-arousal ratings (blue), and activity of different TEM layers (either layer **p**, shown in yellow, or layer **g**, shown in green) for each subject. An exemplary model is shown for hippocampal decoding in subject 1 (S01), although predictive models were trained using BOLD activity separately in each of 5 regions of interest and subject.

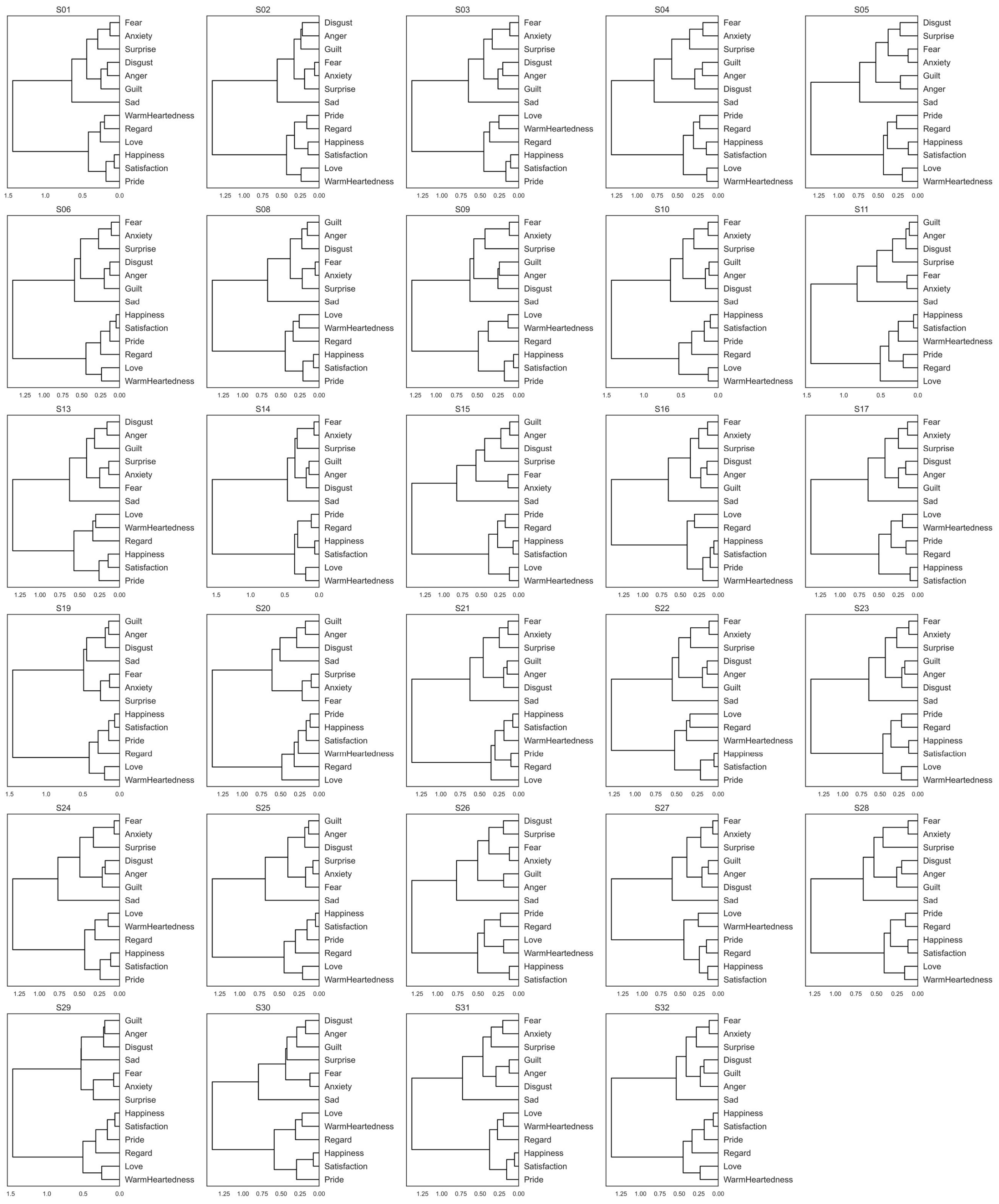


**Supplementary Figure 2**. **Hierarchical clustering of category ratings predicted from hippocampal BOLD activity.** Dendrograms depict the results of agglomerative clustering performed on the pairwise similarity of emotion concepts decoded for each subject. The x-axis indicates the dissimilarity of decoded variables in terms of correlation distance (1 – Pearson's *r*).

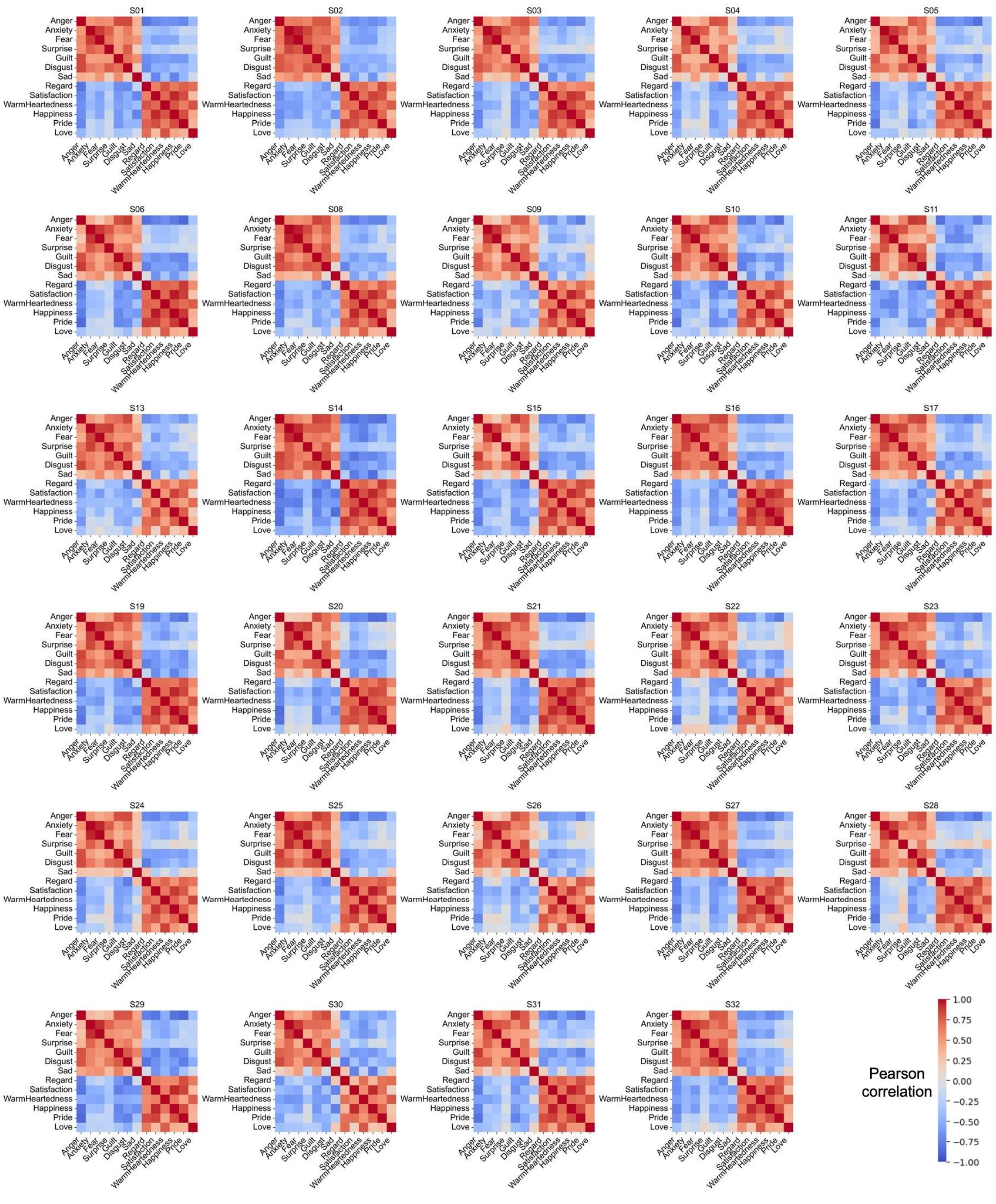


**Supplementary Figure 3**. **Similarity of category ratings predicted from hippocampal BOLD activity.** Correlation matrices (Pearson correlations) depict the pairwise similarity of emotion concepts decoded for each subject. Warm colors indicate positive correlations, and cool colors indicate negative correlations.

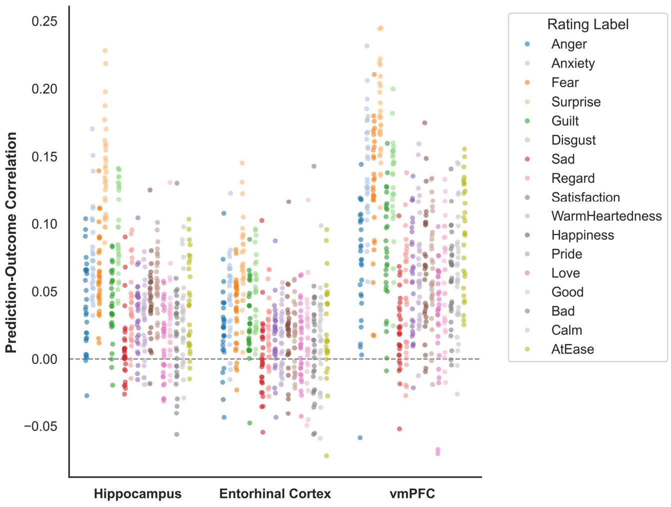


**Supplementary Figure 4**. **Prediction-outcome correlations for individual emotion ratings.** Cross-validated estimates of model performance are shown for each of 13 emotion ratings and 4 dimensional ratings. Each point corresponds to the performance of decoding models in an individual subject in either the hippocampus, entorhinal cortex, or ventromedial prefrontal cortex (vmPFC). The dashed gray line indicates chance.

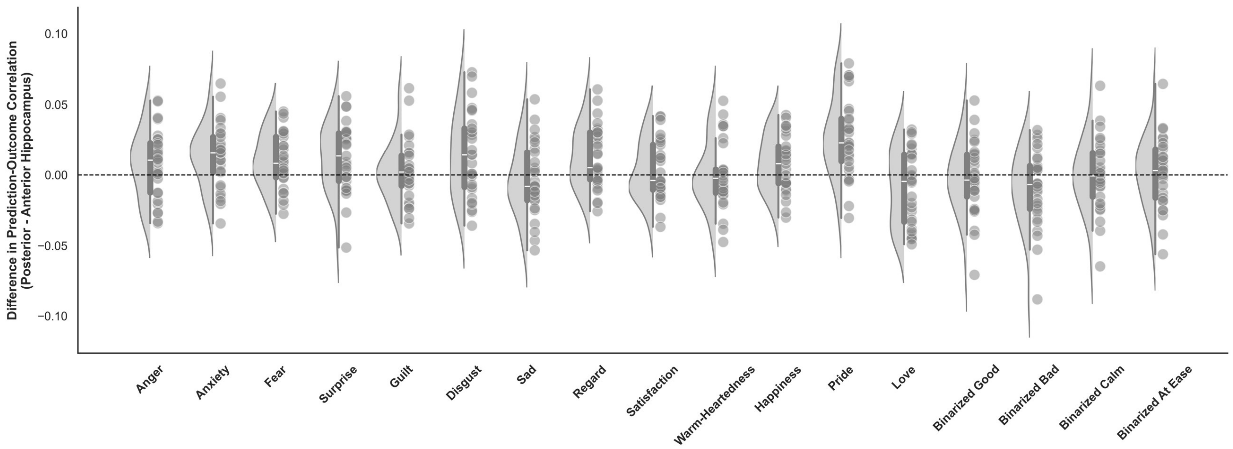


**Supplementary Figure 5**. **Differences of the prediction-outcome correlations between the posterior and anterior hippocampus.** Cross-validated estimates of differences in model performance are shown for each of 13 emotion ratings and 4 binarized valence-arousal ratings. Each point corresponds to the difference in decoding performance of models trained to predict ratings based on BOLD response patterns in posterior and anterior hippocampus. Boxplots indicate the median value and interquartile range. Smooth kernel density estimates are plotted next to individual observations for each rating. The dashed gray line indicates no difference.

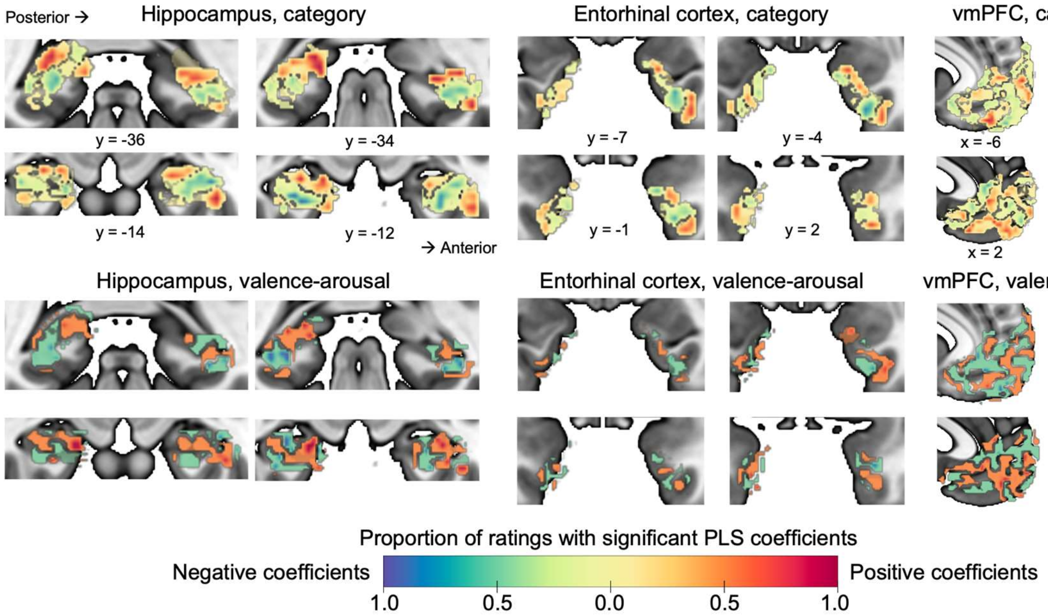


**Supplementary Figure 6**. **Conjunction maps showing regions contributing to the prediction of multiple category ratings and valence-arousal dimensions**. The colormap indicates the proportion of rating items (out of 13 categories or 4 valence-arousal items) with statistically significant PLS coefficients at each voxel (FDR $q < 0.05$). Warm colors denote voxels where coefficients were predominantly positive across items, while cool colors indicate predominantly negative coefficients. vmPFC = ventromedial prefrontal cortex.

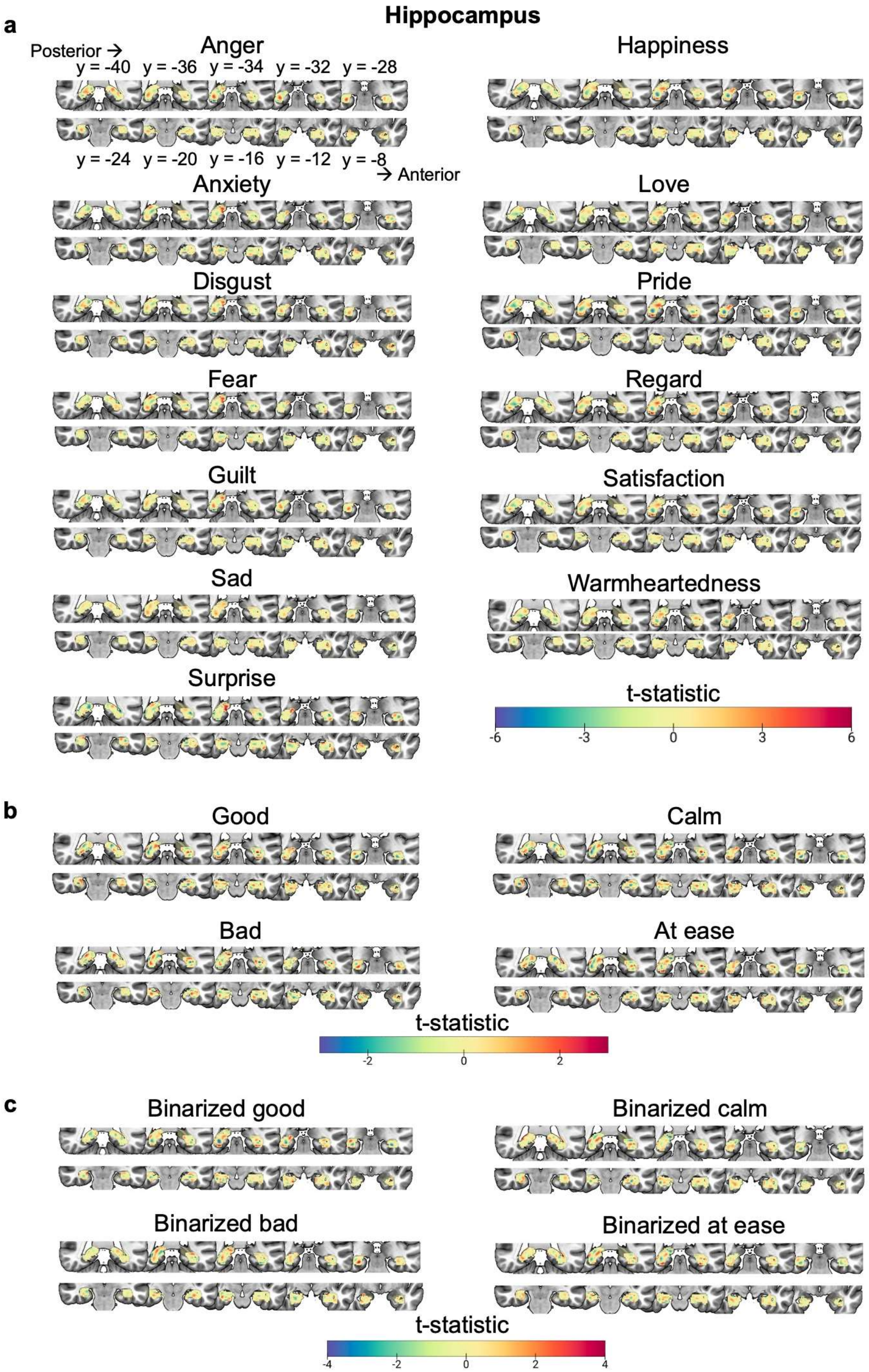


**Supplementary Figure 7**. **Statistical parametric maps of PLS regression coefficients for hippocampal decoders.** Maps reflect the results of one-sample *t*-tests on subject-level PLS coefficients from hippocampal decoders thresholded at FDR $q < 0.05$. Each panel reflects a different decoding model: **(a)** category, **(b)** valence-arousal, and **(c)** binarized valence-arousal.

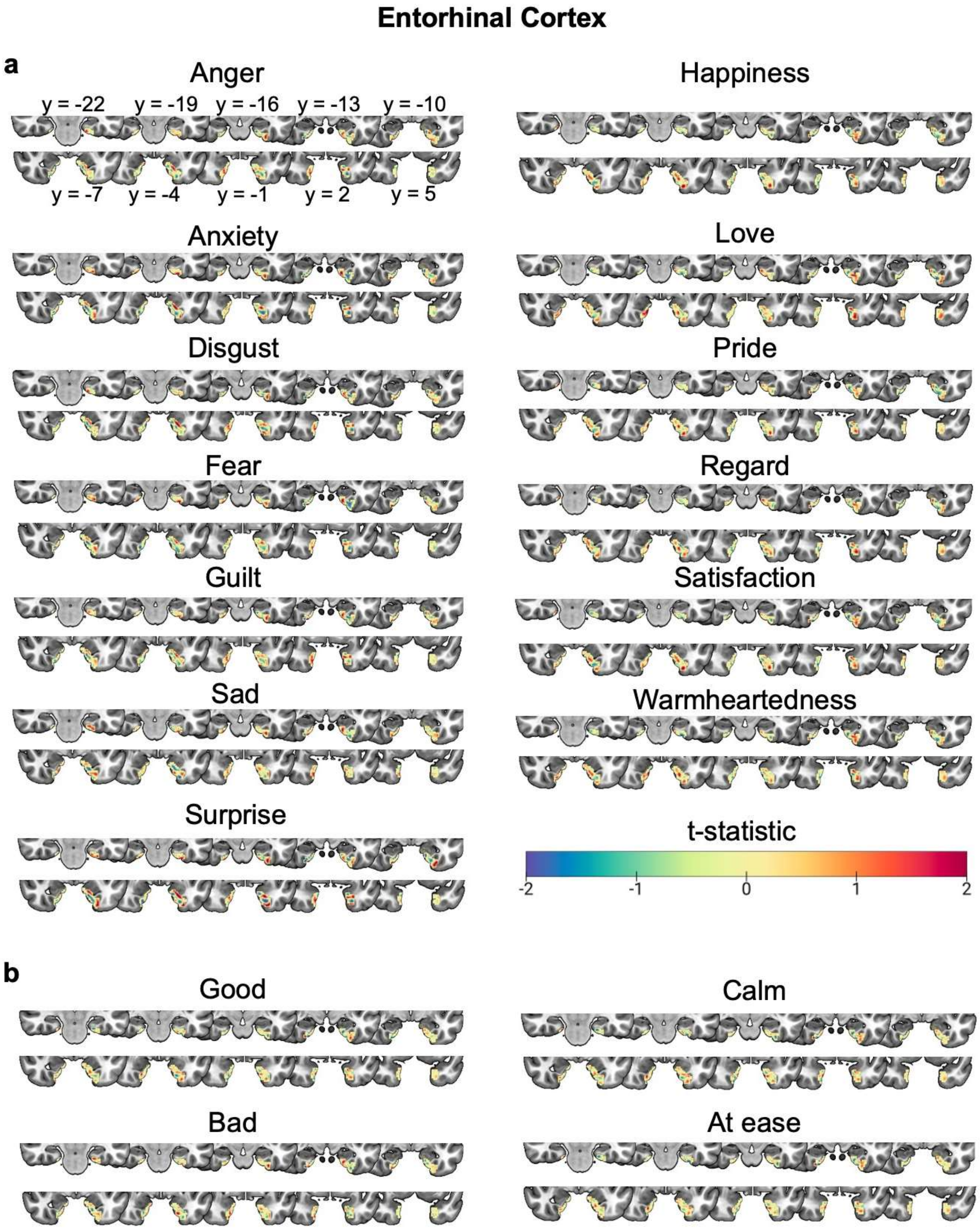


**Supplementary Figure 8**. **Statistical parametric maps of PLS regression coefficients for entorhinal cortex decoders.** Maps reflect the results of one-sample *t*-tests on subject-level PLS coefficients from entorhinal decoders thresholded at FDR $q < 0.05$. Each panel reflects a different decoding model: **(a)** category and **(b)** valence-arousal.

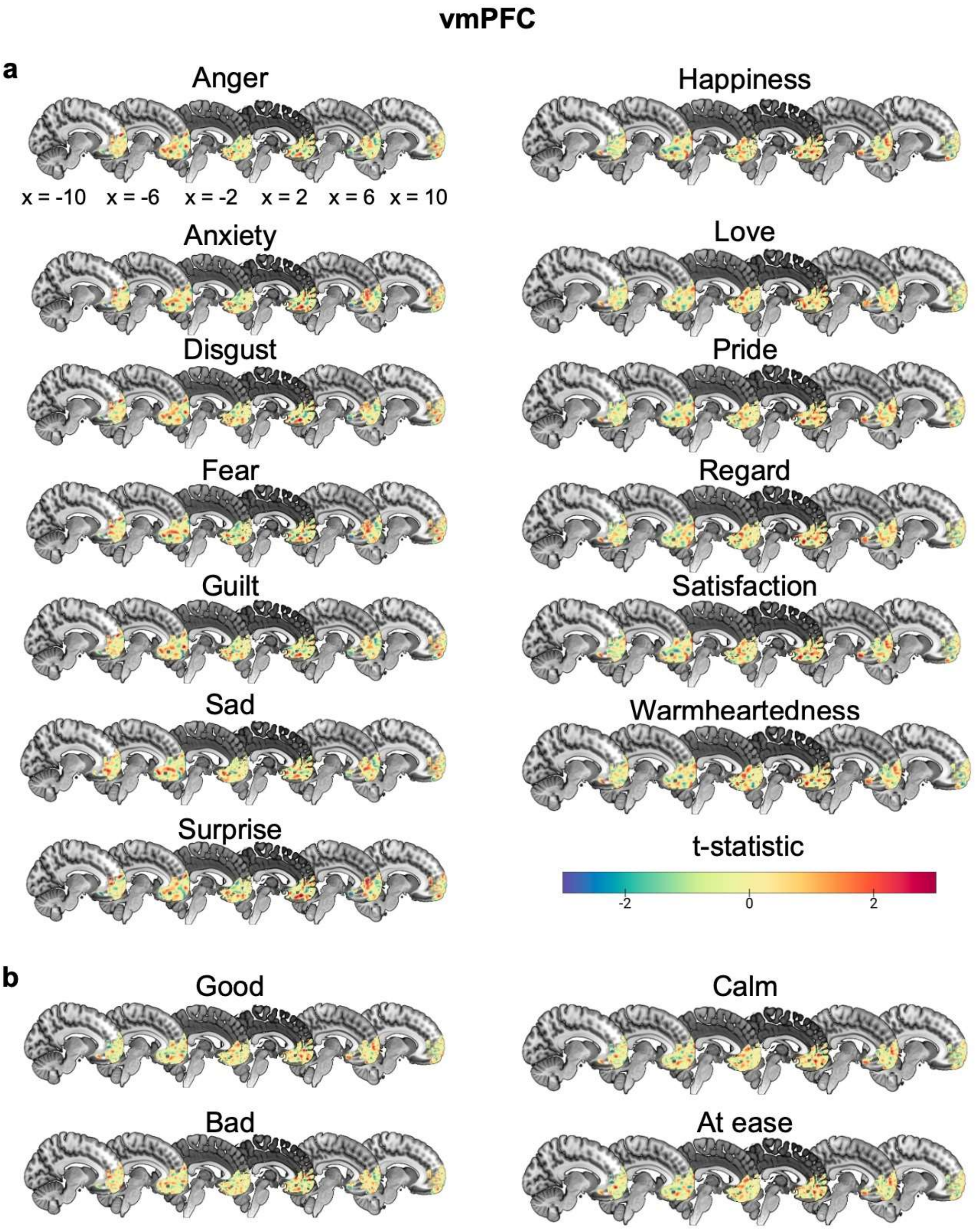


**Supplementary Figure 9**. **Statistical parametric maps of PLS regression coefficients for ventromedial prefrontal cortex (vmPFC) decoders.** Maps reflect the results of one-sample *t*-tests on subject-level PLS coefficients from vmPFC decoders thresholded at FDR $q < 0.05$. Each panel reflects a different decoding model: **(a)** category and **(b)** valence-arousal.

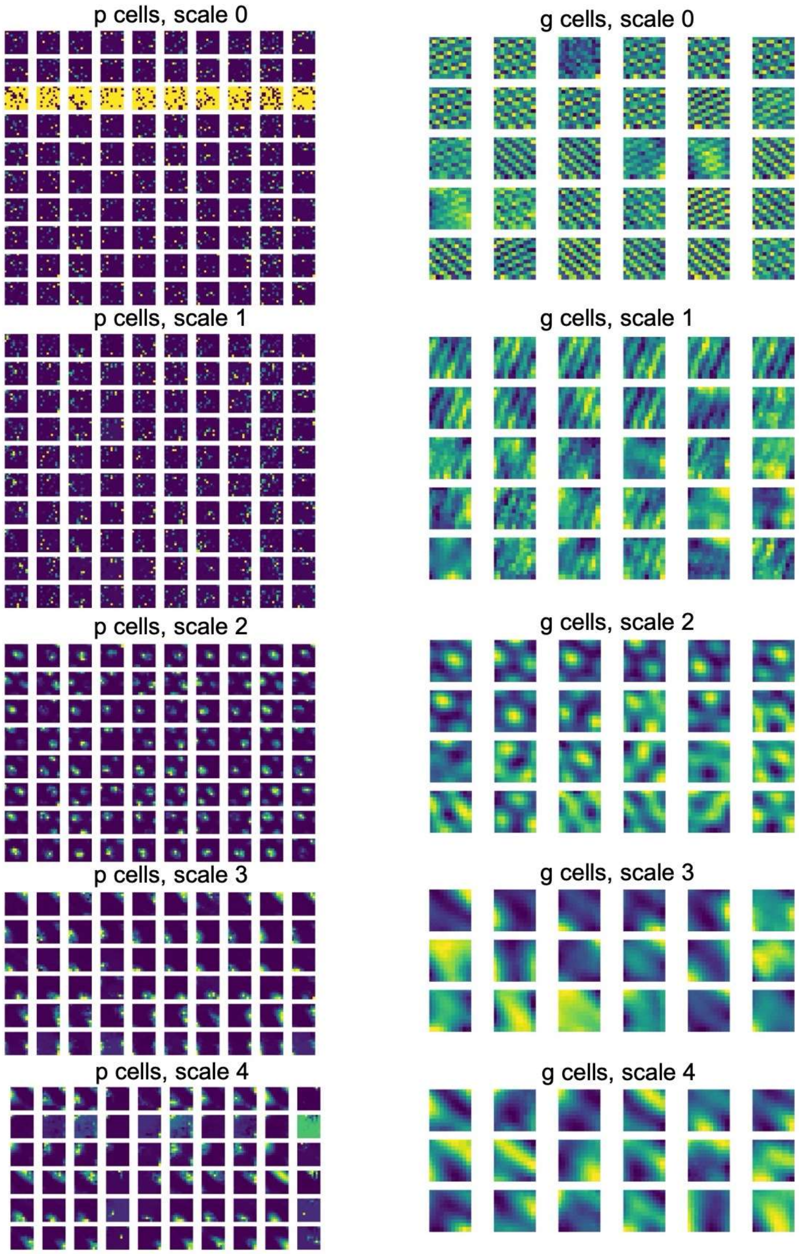


**Supplementary Figure 10**. **Rate maps of units in layers p and g of the Tolman Eichenbaum Machine.** Rows organize units based on representational scale.

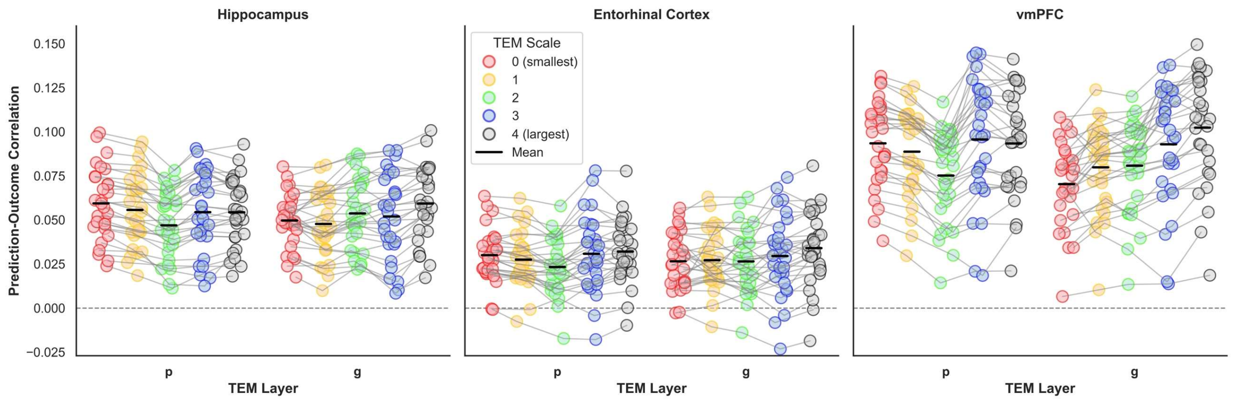


**Supplementary Figure 11**. **Performance of decoding models trained to predict activity of p and g cells from BOLD activity in hippocampus, entorhinal cortex, and ventromedial prefrontal cortex (vmPFC).** Each point represents the prediction-outcome correlation averaged across units at each scale for one subject. Horizontal black lines indicate the group average, connected gray lines indicate performance for one subject, and the horizontal dashed line indicates chance. TEM = Tolman Eichenbaum Machine.

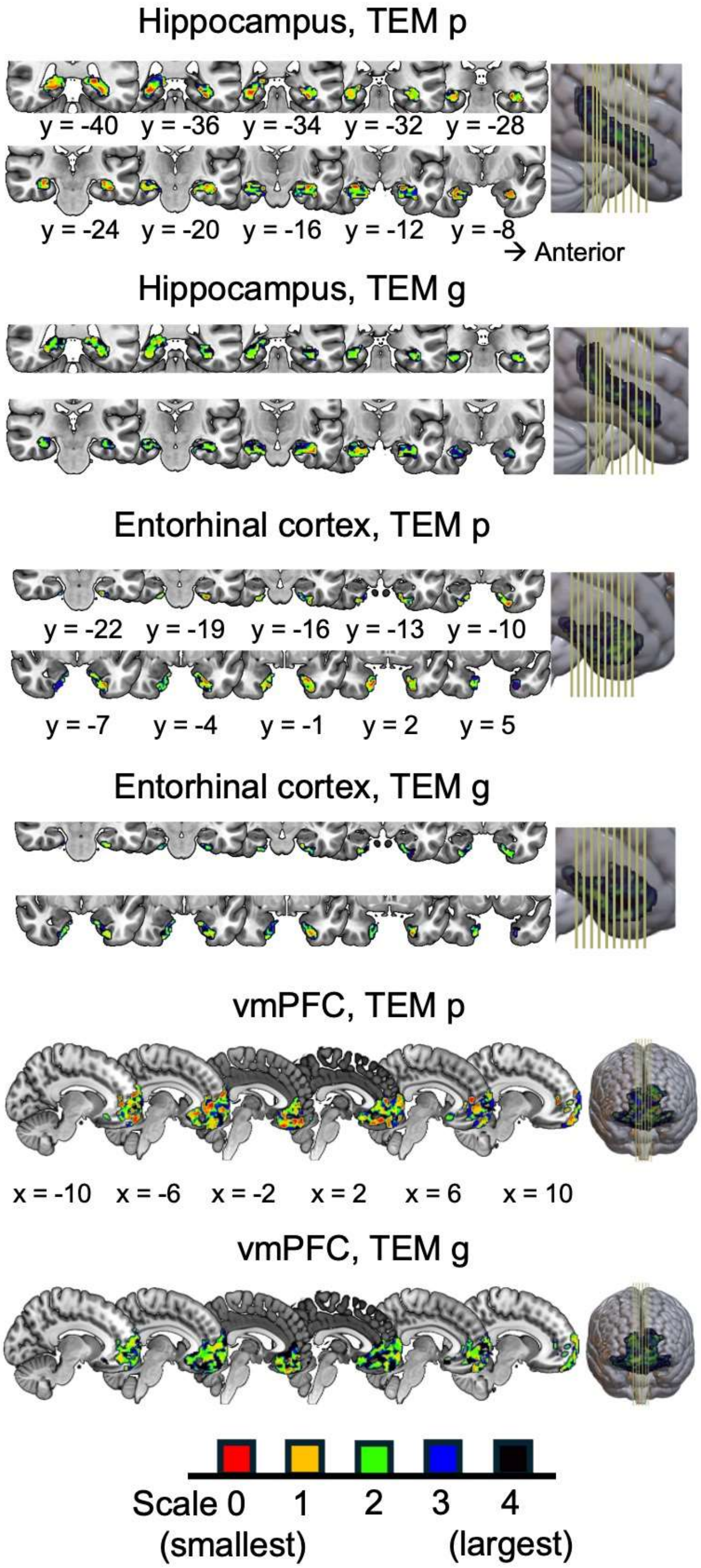


**Supplementary Figure 12**. Maximum representational scale maps for the hippocampus, entorhinal cortex, and ventromedial prefrontal cortex (vmPFC). Maps show the representational scale with the largest PLS coefficient. Warm colors indicate the presence of small-scale representations and cool colors indicate larger scales. On the right, brain renders are shown with yellow lines indicating the locations of the slices displayed on the left.

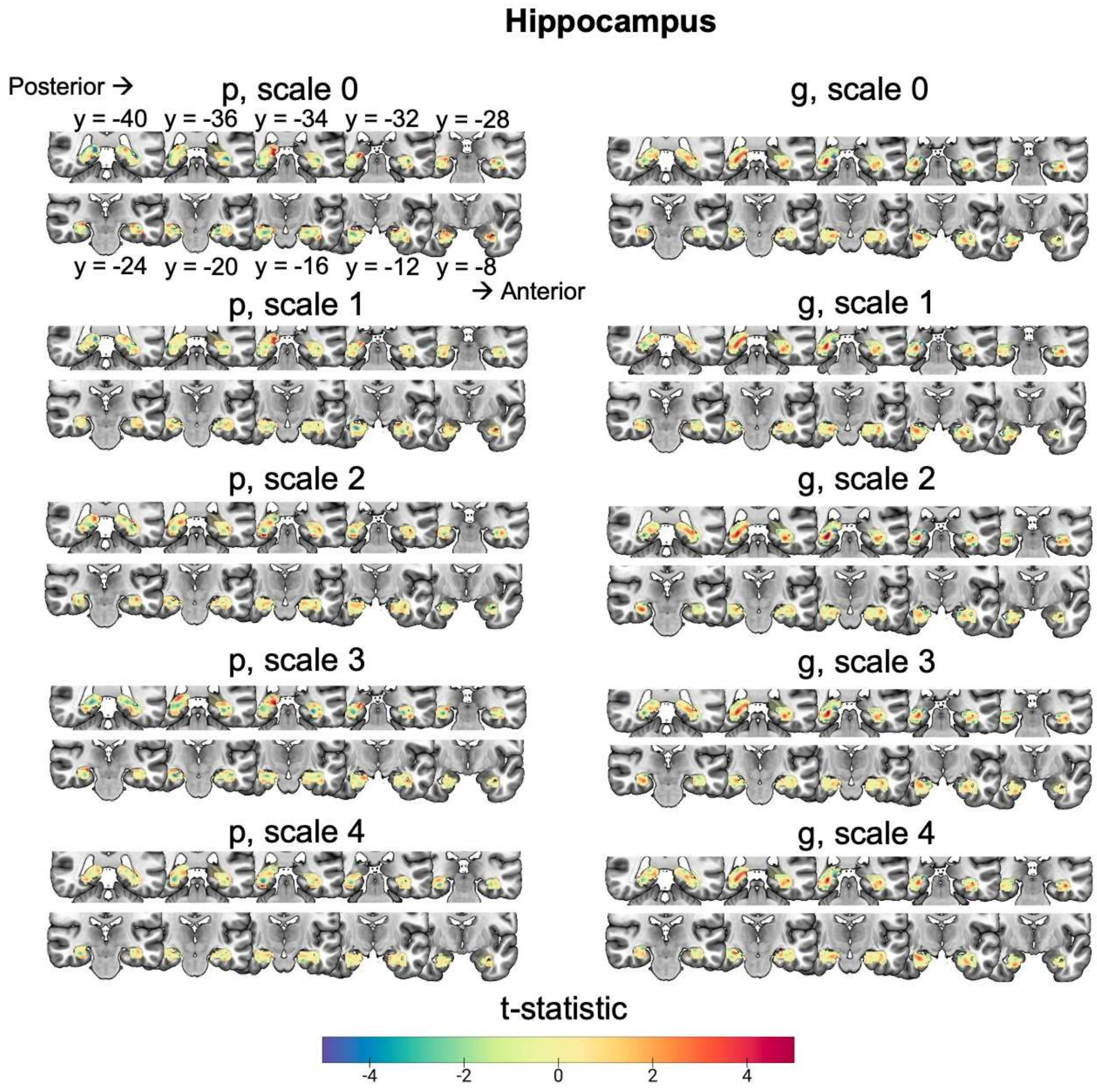


**Supplementary Figure 13**. t-statistic maps (thresholded at FDR $q < 0.05$) derived from group-level inference on hippocampal PLS coefficient maps averaged across units for each scale of each TEM layer.

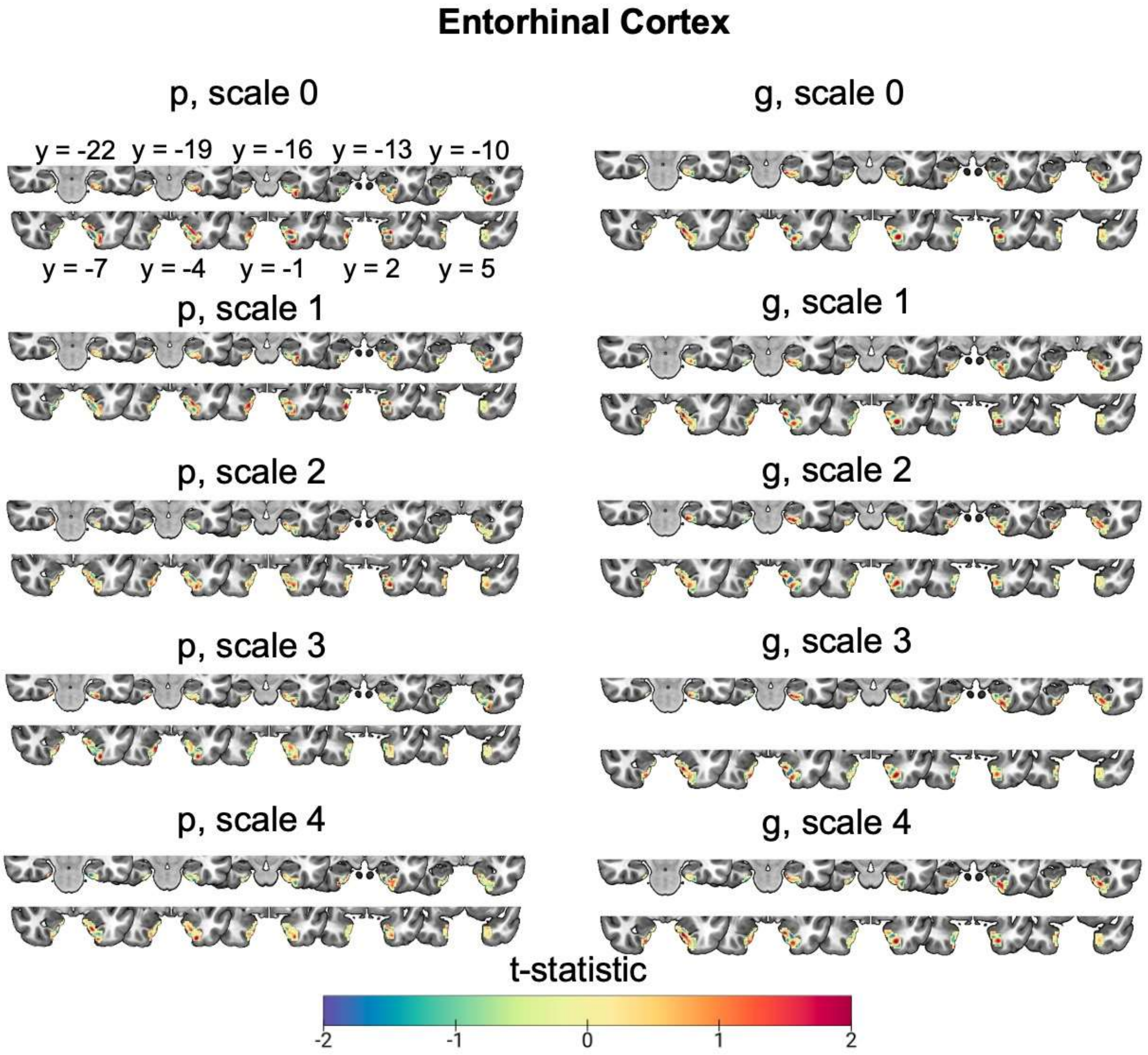


**Supplementary Figure 14**. t-statistic maps (thresholded at FDR $q < 0.05$) derived from group-level inference on entorhinal PLS coefficient maps averaged across units for each scale of each TEM layer.

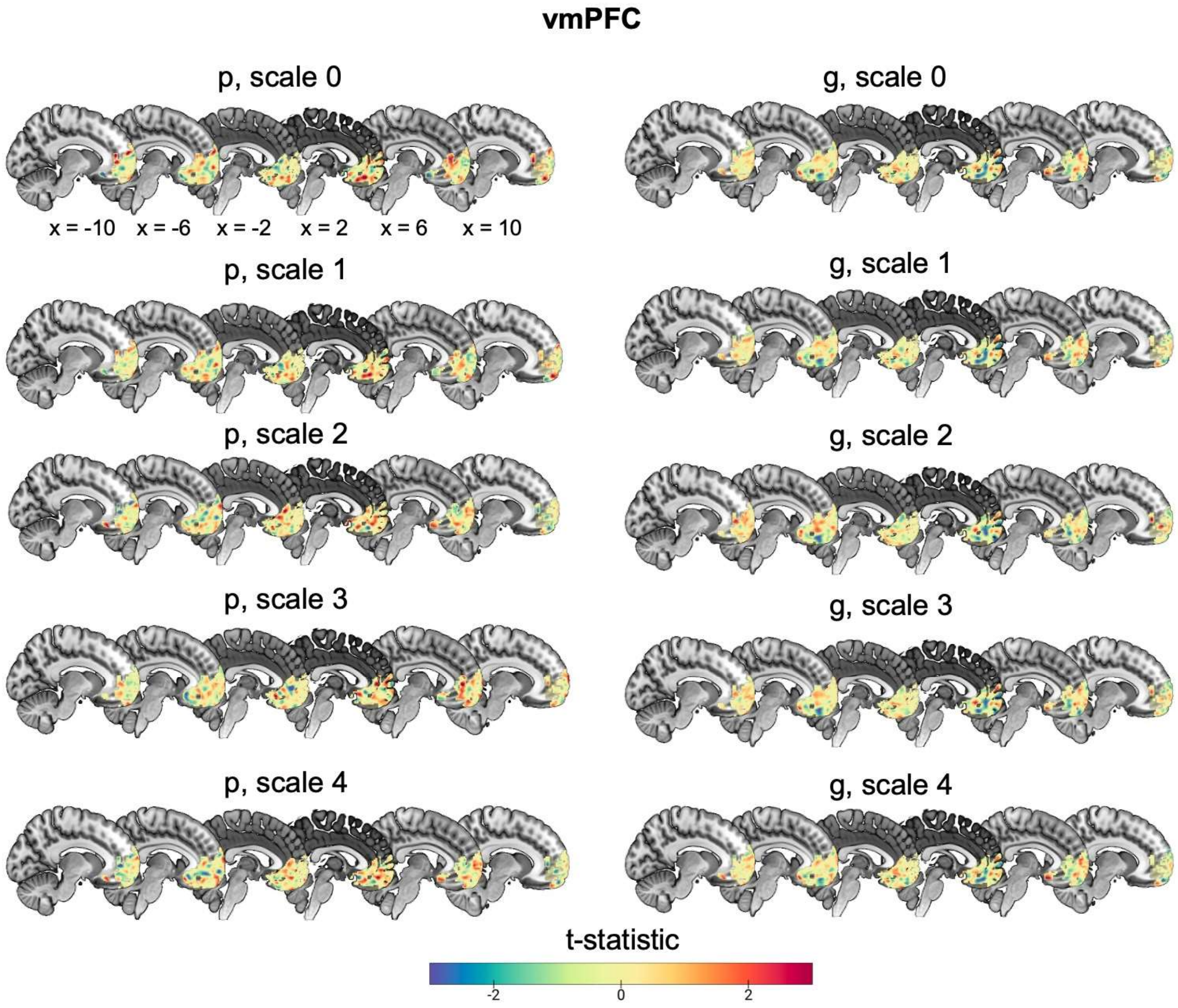


**Supplementary Figure 15**. t-statistic maps (thresholded at FDR $q < 0.05$) derived from group-level inference on vmPFC PLS coefficient maps averaged across units for each scale of each TEM layer.

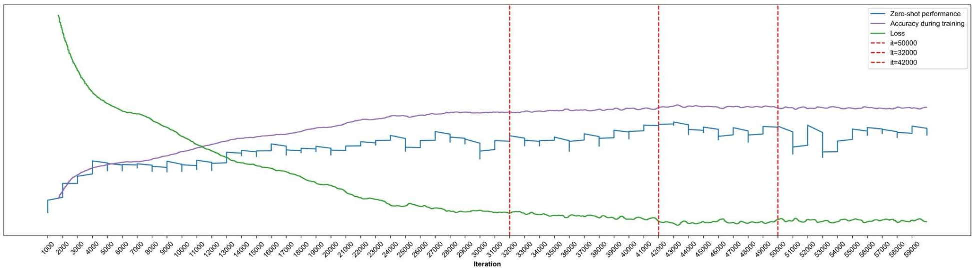


**Supplementary Figure 16.** Loss and performance across iterations of TEM. The 'Accuracy during training' curve shows the accuracy of predicting the next sensory observation during training. The 'Zero-shot performance' curve shows the accuracy of predicting observations following a new action to a known location in the testing environment. The 'Loss' curve shows the training loss across iterations. Vertical red lines indicate the iterations used in our analyses.

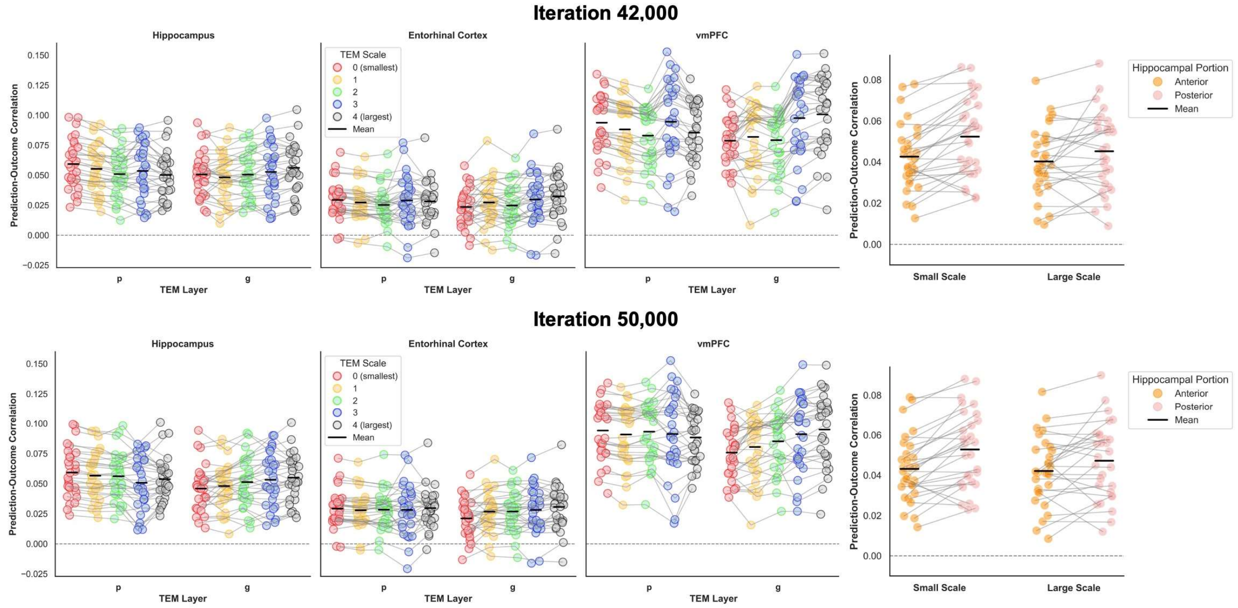


**Supplementary Figure 17**. Results from different iterations of TEM (see Fig. 4b in the main text and Supplementary Fig. 11).

### Supplementary Tables

**Supplementary Table 1**. **One-sample *t*-tests ($H_0$: $\mu = 0$) for Fisher z-transformed prediction-outcome correlations in decoding emotion ratings from BOLD activity**

| Region | Emotion | *t* | *p* | FDR *q* |
|---|---|---|---|---|
| Hippocampus | Anger | 6.46 | < 0.0001 | < 0.0001 |
| Entorhinal cortex | Anger | 4.02 | 0.0002 | 0.0003 |
| vmPFC | Anger | 8.44 | < 0.0001 | < 0.0001 |
| Hippocampus | Anxiety | 12.55 | < 0.0001 | < 0.0001 |
| Entorhinal cortex | Anxiety | 8.07 | < 0.0001 | < 0.0001 |
| vmPFC | Anxiety | 19.04 | < 0.0001 | < 0.0001 |
| Hippocampus | Fear | 11.62 | < 0.0001 | < 0.0001 |
| Entorhinal cortex | Fear | 7.13 | < 0.0001 | < 0.0001 |
| vmPFC | Fear | 14.46 | < 0.0001 | < 0.0001 |
| Hippocampus | Surprise | 20.90 | < 0.0001 | < 0.0001 |
| Entorhinal cortex | Surprise | 12.66 | < 0.0001 | < 0.0001 |
| vmPFC | Surprise | 29.42 | < 0.0001 | < 0.0001 |
| Hippocampus | Guilt | 8.25 | < 0.0001 | < 0.0001 |
| Entorhinal cortex | Guilt | 5.71 | < 0.0001 | < 0.0001 |
| vmPFC | Guilt | 10.17 | < 0.0001 | < 0.0001 |
| Hippocampus | Disgust | 15.64 | < 0.0001 | < 0.0001 |
| Entorhinal cortex | Disgust | 9.59 | < 0.0001 | < 0.0001 |
| vmPFC | Disgust | 18.73 | < 0.0001 | < 0.0001 |
| Hippocampus | Sad | 1.91 | 0.0329 | 0.0343 |
| Entorhinal cortex | Sad | 0.18 | 0.4281 | 0.4281 |
| vmPFC | Sad | 3.94 | 0.0002 | 0.0003 |
| Hippocampus | Regard | 7.15 | < 0.0001 | < 0.0001 |
| Entorhinal cortex | Regard | 3.78 | 0.0004 | 0.0005 |
| vmPFC | Regard | 5.83 | < 0.0001 | < 0.0001 |
| Hippocampus | Satisfaction | 6.92 | < 0.0001 | < 0.0001 |
| Entorhinal cortex | Satisfaction | 3.50 | 0.0008 | 0.0010 |
| vmPFC | Satisfaction | 7.89 | < 0.0001 | < 0.0001 |
| Hippocampus | WarmHeartedness | 6.00 | < 0.0001 | < 0.0001 |
| Entorhinal cortex | WarmHeartedness | 3.51 | 0.0008 | 0.0009 |
| vmPFC | WarmHeartedness | 8.86 | < 0.0001 | < 0.0001 |
| Hippocampus | Happiness | 8.41 | < 0.0001 | < 0.0001 |
| Entorhinal cortex | Happiness | 3.63 | 0.0006 | 0.0007 |
| vmPFC | Happiness | 7.67 | < 0.0001 | < 0.0001 |
| Hippocampus | Pride | 13.06 | < 0.0001 | < 0.0001 |

| | | | | |
|---|---|---|---|---|
| Entorhinal cortex | Pride | 4.03 | 0.0002 | 0.0003 |
| vmPFC | Pride | 8.99 | < 0.0001 | < 0.0001 |
| Hippocampus | Love | 2.44 | 0.0106 | 0.0117 |
| Entorhinal cortex | Love | 2.31 | 0.0142 | 0.0154 |
| vmPFC | Love | 4.46 | 0.0001 | 0.0001 |
| Hippocampus | Good | 5.22 | < 0.0001 | < 0.0001 |
| Entorhinal cortex | Good | 2.47 | 0.0099 | 0.0112 |
| vmPFC | Good | 7.78 | < 0.0001 | < 0.0001 |
| Hippocampus | Bad | 2.29 | 0.0148 | 0.0157 |
| Entorhinal cortex | Bad | 1.43 | 0.0821 | 0.0838 |
| vmPFC | Bad | 9.20 | < 0.0001 | < 0.0001 |
| Hippocampus | Calm | 5.18 | < 0.0001 | < 0.0001 |
| Entorhinal cortex | Calm | 2.77 | 0.0049 | 0.0057 |
| vmPFC | Calm | 7.89 | < 0.0001 | < 0.0001 |
| Hippocampus | AtEase | 6.41 | < 0.0001 | < 0.0001 |
| Entorhinal cortex | AtEase | 3.39 | 0.0011 | 0.0013 |
| vmPFC | AtEase | 12.45 | < 0.0001 | < 0.0001 |

**Supplementary Table 2**. **Paired *t*-tests for testing whether the Fisher z-transformed prediction-outcome correlation is higher in posterior than anterior hippocampus for each emotion category**

| | *t* | *p* | FDR *q* |
|---|---|---|---|
| Regard | 3.06 | 0.0024 | 0.0106 |
| Satisfaction | 0.80 | 0.2150 | 0.3105 |
| WarmHeartedness | 0.08 | 0.4688 | 0.5541 |
| Happiness | 2.03 | 0.0259 | 0.0481 |
| Pride | 5.15 | < 0.0001 | 0.0001 |
| Love | -1.76 | 0.9554 | 0.9554 |
| Anger | 1.72 | 0.0486 | 0.0789 |
| Anxiety | 3.22 | 0.0016 | 0.0106 |
| Fear | 2.79 | 0.0046 | 0.0124 |
| Guilt | 0.60 | 0.2778 | 0.3611 |
| Disgust | 2.59 | 0.0075 | 0.0163 |
| Sad | -0.66 | 0.7441 | 0.8061 |
| Surprise | 2.78 | 0.0048 | 0.0124 |

**Supplementary Table 3. One-sample *t*-tests ($H_0$: $\mu = 0$) for Fisher z-transformed prediction-outcome correlations in decoding TEM activity at iteration 32,000 from BOLD activity**

| Region | layer | scale | *t* | *p* | FDR *q* |
|---|---|---|---|---|---|
| Entorhinal cortex | g | 0 | 37.33 | < 0.0001 | < 0.0001 |
| Entorhinal cortex | g | 1 | 32.58 | < 0.0001 | < 0.0001 |
| Entorhinal cortex | g | 2 | 23.51 | < 0.0001 | < 0.0001 |
| Entorhinal cortex | g | 3 | 23.71 | < 0.0001 | < 0.0001 |
| Entorhinal cortex | g | 4 | 26.77 | < 0.0001 | < 0.0001 |
| Entorhinal cortex | p | 0 | 54.78 | < 0.0001 | < 0.0001 |
| Entorhinal cortex | p | 1 | 56.96 | < 0.0001 | < 0.0001 |
| Entorhinal cortex | p | 2 | 40.53 | < 0.0001 | < 0.0001 |
| Entorhinal cortex | p | 3 | 45.76 | < 0.0001 | < 0.0001 |
| Entorhinal cortex | p | 4 | 59.24 | < 0.0001 | < 0.0001 |
| Hippocampus | g | 0 | 67.23 | < 0.0001 | < 0.0001 |
| Hippocampus | g | 1 | 51.41 | < 0.0001 | < 0.0001 |
| Hippocampus | g | 2 | 43.55 | < 0.0001 | < 0.0001 |
| Hippocampus | g | 3 | 37.26 | < 0.0001 | < 0.0001 |
| Hippocampus | g | 4 | 43.78 | < 0.0001 | < 0.0001 |
| Hippocampus | p | 0 | 75.43 | < 0.0001 | < 0.0001 |
| Hippocampus | p | 1 | 82.20 | < 0.0001 | < 0.0001 |
| Hippocampus | p | 2 | 64.55 | < 0.0001 | < 0.0001 |
| Hippocampus | p | 3 | 65.35 | < 0.0001 | < 0.0001 |
| Hippocampus | p | 4 | 87.62 | < 0.0001 | < 0.0001 |
| vmPFC | g | 0 | 63.65 | < 0.0001 | < 0.0001 |
| vmPFC | g | 1 | 56.89 | < 0.0001 | < 0.0001 |
| vmPFC | g | 2 | 42.26 | < 0.0001 | < 0.0001 |
| vmPFC | g | 3 | 49.73 | < 0.0001 | < 0.0001 |
| vmPFC | g | 4 | 52.90 | < 0.0001 | < 0.0001 |
| vmPFC | p | 0 | 101.61 | < 0.0001 | < 0.0001 |
| vmPFC | p | 1 | 108.15 | < 0.0001 | < 0.0001 |
| vmPFC | p | 2 | 80.83 | < 0.0001 | < 0.0001 |
| vmPFC | p | 3 | 88.23 | < 0.0001 | < 0.0001 |
| vmPFC | p | 4 | 106.30 | < 0.0001 | < 0.0001 |

**Supplementary Table 4. One-sample *t*-tests ($H_0$: $\mu = 0$) for Fisher z-transformed prediction-outcome correlations in decoding TEM activity at iteration 42,000 from BOLD activity**

| Region | TEM layer | Scale | *t* | *p* (one-tailed) | FDR *q* |
|---|---|---|---|---|---|
| Entorhinal cortex | g | 0 | 32.27 | < 0.0001 | < 0.0001 |
| Entorhinal cortex | g | 1 | 30.11 | < 0.0001 | < 0.0001 |
| Entorhinal cortex | g | 2 | 22.26 | < 0.0001 | < 0.0001 |
| Entorhinal cortex | g | 3 | 23.39 | < 0.0001 | < 0.0001 |
| Entorhinal cortex | g | 4 | 25.67 | < 0.0001 | < 0.0001 |
| Entorhinal cortex | p | 0 | 54.92 | < 0.0001 | < 0.0001 |
| Entorhinal cortex | p | 1 | 52.65 | < 0.0001 | < 0.0001 |
| Entorhinal cortex | p | 2 | 39.76 | < 0.0001 | < 0.0001 |
| Entorhinal cortex | p | 3 | 42.06 | < 0.0001 | < 0.0001 |
| Entorhinal cortex | p | 4 | 47.16 | < 0.0001 | < 0.0001 |
| Hippocampus | g | 0 | 60.65 | < 0.0001 | < 0.0001 |
| Hippocampus | g | 1 | 48.45 | < 0.0001 | < 0.0001 |
| Hippocampus | g | 2 | 38.20 | < 0.0001 | < 0.0001 |
| Hippocampus | g | 3 | 34.64 | < 0.0001 | < 0.0001 |
| Hippocampus | g | 4 | 41.07 | < 0.0001 | < 0.0001 |
| Hippocampus | p | 0 | 75.77 | < 0.0001 | < 0.0001 |
| Hippocampus | p | 1 | 75.27 | < 0.0001 | < 0.0001 |
| Hippocampus | p | 2 | 58.47 | < 0.0001 | < 0.0001 |
| Hippocampus | p | 3 | 64.40 | < 0.0001 | < 0.0001 |
| Hippocampus | p | 4 | 69.37 | < 0.0001 | < 0.0001 |
| vmPFC | g | 0 | 69.39 | < 0.0001 | < 0.0001 |
| vmPFC | g | 1 | 56.88 | < 0.0001 | < 0.0001 |
| vmPFC | g | 2 | 41.76 | < 0.0001 | < 0.0001 |
| vmPFC | g | 3 | 49.44 | < 0.0001 | < 0.0001 |
| vmPFC | g | 4 | 50.99 | < 0.0001 | < 0.0001 |
| vmPFC | p | 0 | 105.87 | < 0.0001 | < 0.0001 |
| vmPFC | p | 1 | 99.08 | < 0.0001 | < 0.0001 |
| vmPFC | p | 2 | 77.25 | < 0.0001 | < 0.0001 |
| vmPFC | p | 3 | 86.31 | < 0.0001 | < 0.0001 |
| vmPFC | p | 4 | 85.77 | < 0.0001 | < 0.0001 |

**Supplementary Table 5. One-sample *t*-tests ($H_0$: $\mu = 0$) for Fisher z-transformed prediction-outcome correlations in decoding TEM activity at iteration 50,000 from BOLD activity**

| Region | TEM layer | Scale | *t* | *p* (one-tailed) | FDR *q* |
|---|---|---|---|---|---|
| Entorhinal cortex | g | 0 | 30.33 | < 0.0001 | < 0.0001 |
| Entorhinal cortex | g | 1 | 30.92 | < 0.0001 | < 0.0001 |
| Entorhinal cortex | g | 2 | 23.91 | < 0.0001 | < 0.0001 |
| Entorhinal cortex | g | 3 | 21.70 | < 0.0001 | < 0.0001 |
| Entorhinal cortex | g | 4 | 25.22 | < 0.0001 | < 0.0001 |
| Entorhinal cortex | p | 0 | 55.07 | < 0.0001 | < 0.0001 |
| Entorhinal cortex | p | 1 | 54.87 | < 0.0001 | < 0.0001 |
| Entorhinal cortex | p | 2 | 49.41 | < 0.0001 | < 0.0001 |
| Entorhinal cortex | p | 3 | 43.13 | < 0.0001 | < 0.0001 |
| Entorhinal cortex | p | 4 | 42.91 | < 0.0001 | < 0.0001 |
| Hippocampus | g | 0 | 55.71 | < 0.0001 | < 0.0001 |
| Hippocampus | g | 1 | 49.18 | < 0.0001 | < 0.0001 |
| Hippocampus | g | 2 | 39.62 | < 0.0001 | < 0.0001 |
| Hippocampus | g | 3 | 36.96 | < 0.0001 | < 0.0001 |
| Hippocampus | g | 4 | 38.90 | < 0.0001 | < 0.0001 |
| Hippocampus | p | 0 | 76.00 | < 0.0001 | < 0.0001 |
| Hippocampus | p | 1 | 78.40 | < 0.0001 | < 0.0001 |
| Hippocampus | p | 2 | 72.72 | < 0.0001 | < 0.0001 |
| Hippocampus | p | 3 | 67.35 | < 0.0001 | < 0.0001 |
| Hippocampus | p | 4 | 55.17 | < 0.0001 | < 0.0001 |
| vmPFC | g | 0 | 69.03 | < 0.0001 | < 0.0001 |
| vmPFC | g | 1 | 58.89 | < 0.0001 | < 0.0001 |
| vmPFC | g | 2 | 45.43 | < 0.0001 | < 0.0001 |
| vmPFC | g | 3 | 44.59 | < 0.0001 | < 0.0001 |
| vmPFC | g | 4 | 47.35 | < 0.0001 | < 0.0001 |
| vmPFC | p | 0 | 106.69 | < 0.0001 | < 0.0001 |
| vmPFC | p | 1 | 105.38 | < 0.0001 | < 0.0001 |
| vmPFC | p | 2 | 93.77 | < 0.0001 | < 0.0001 |
| vmPFC | p | 3 | 85.22 | < 0.0001 | < 0.0001 |
| vmPFC | p | 4 | 78.98 | < 0.0001 | < 0.0001 |

**Supplementary Table 6. Results from different iterations of TEM.**

| Analysis | Iteration 32,000 | Iteration 42,000 | Iteration 50,000 |
|---|---|---|---|
| **Hippocampus Only** | | | |
| **p** vs. **g** | Δ*z* = 0.0017, 95% bootstrap *CI* [0.0001, 0.0034], $p = 0.0145$ | Δ*z* = 0.0023, 95% bootstrap *CI* [0.0008, 0.0038], $p = 0.0062$ | Δ*z* = 0.0047, 95% bootstrap *CI* [0.0031, 0.0062], $p < .0001$ |
| Linear contrast of scale: **p** vs. **g** | Δ*z* = -0.0352, 95% bootstrap *CI* [-0.0448, -0.0254], $p < .0001$ | Δ*z* = -0.0358, 95% bootstrap *CI* [-0.0447, -0.0268], $p < .0001$ | Δ*z* = -0.0407, 95% bootstrap *CI* [-0.0496, -0.0317], $p < .0001$ |
| Category vs. valence-arousal × correlation vs. partial correlation | Δz = 0.0039, 95% bootstrap *CI* [0.0021, 0.0057], $p = .0010$ | Δz = 0.0037, 95% bootstrap *CI* [0.0018, 0.0055], $p = 0.0010$ | Δ*z* = 0.0038, 95% bootstrap *CI* [0.0019, 0.0057], $p = 0.0016$ |
| Posterior vs. anterior hippocampus × small vs. large scale | Δ*z* = 0.0052, 95% bootstrap *CI* [0.0033, 0.0071], $p = .0019$ | Δ*z* = 0.0053, 95% bootstrap CI [0.0033, 0.0072], $p = .0010$ | Δ*z* = 0.0049, 95% bootstrap CI [0.0029, 0.0069], $p = .0015$ |
| **Hippocampus vs. Entorhinal Cortex vs. vmPFC** | | | |
| Region × TEM layer × scale | $F(8, 616) = 12.44$, $p < .0001$ | $F(8, 616) = 9.24$, $p < .0001$ | $F(8, 616) = 6.40$, $p < .0001$ |
| *Linear Contrast of Scale:* | | | |
| Entorhinal cortex: **p** vs. **g** | Δz = -0.0099, 95% bootstrap *CI* [-0.0199, -0.0001], $p = .0461$ | Δz = -0.0209, 95% bootstrap *CI* [-0.0301, -0.0119], $p = .0006$ | Δz = -0.0194, 95% bootstrap *CI* [-0.0286, -0.0104], $p = 0.0009$ |
| vmPFC: **p** vs. **g** | Δ*z* = -0.0712, 95% bootstrap *CI* [-0.0809, -0.0614], $p < .0001$ | Δ*z* = -0.0706, 95% bootstrap *CI* [-0.0796, -0.0616], $p < .0001$ | Δ*z* = -0.0608, 95% bootstrap *CI* [-0.0697, -0.0518], $p < .0001$ |
| **p**: hippocampus vs. entorhinal cortex | Δz = -0.0191, 95% bootstrap *CI* [-0.0289, -0.0093], $p = .0003$ | Δz = -0.0196, 95% bootstrap *CI* [-0.0287, -0.0107], $p < .0001$ | Δz = -0.0182, 95% bootstrap *CI* [-0.0273, -0.0093], $p < .0001$ |
| **p**: hippocampus vs. vmPFC | Δz = -0.0182, 95% bootstrap *CI* [-0.0279, -0.0086], $p = .0031$ | Δz = -0.0097, 95% bootstrap *CI* [-0.0187, -0.0008], $p = 0.0095$ | Δz = -0.0059, 95% bootstrap *CI* [-0.0149, 0.0029], $p = 0.0603$ |
| **g**: hippocampus vs. vmPFC | Δz = -0.0542, 95% bootstrap *CI* [-0.0640, -0.0447], $p < .0001$ | Δz = -0.0445, 95% bootstrap *CI* [-0.0535, -0.0358], $p < .0001$ | Δz = -0.0261, 95% bootstrap *CI* [-0.0350, -0.0173], $p < .0001$ |
| **g**: entorhinal cortex vs. vmPFC | Δz = -0.0604, 95% bootstrap *CI* [-0.0700, -0.0506], $p < .0001$ | Δz = -0.0398, 95% bootstrap *CI* [-0.0486, -0.0307], $p < .0001$ | Δz = -0.0291, 95% bootstrap *CI* [-0.0378, -0.0200], $p = .0005$ |
